# An integrated multi-omic atlas of the choroid plexus highlights conserved molecular programs of epithelial function

**DOI:** 10.64898/2026.10.05.756688

**Authors:** Evan Dennis, Ethan Tieu, Stuart Benjamin Fass, Lorenzo F. Ochoa, Qiang Li, Nicolas Pigadas, William C. Davalan, Martin Kahn, Kyriakitsa Galani, Phan Q. Duy, Baojian Fan, Neel H. Mehta, Hector W. Cure, Milica Nenadovich, Seth L. Alper, Kwanghun Chung, Li-Huei Tsai, Manolis Kellis, Kristopher T. Kahle

**Affiliations:** Department of Neurosurgery, Massachusetts General Hospital & Harvard Medical School, Boston, MA 02114, USA; Broad Institute of Harvard and Massachusetts Institute of Technology, Cambridge, MA 02142, USA; Computer Science and Artificial Intelligence Laboratory, Massachusetts Institute of Technology, Cambridge, MA, 02139, USA; Picower Institute for Learning and Memory, Massachusetts Institute of Technology, Cambridge, MA 02139, USA; Department of Brain and Cognitive Sciences, Massachusetts Institute of Technology, Cambridge, MA 02139, USA; Division of Neurosurgery, Department of Surgery, University of Alberta, Edmonton, AB T6G 2R3, CA; Division of Nephrology and Center for Vascular Biology Research, Beth Israel Deaconess Medical Center & Harvard Medical School, Boston, MA 02215, USA; Institute for Medical Engineering and Science, Massachusetts Institute of Technology, Cambridge, MA, USA; Department of Chemical Engineering, Massachusetts Institute of Technology, Cambridge, MA, USA; Program in Neuroscience (PiN), Harvard University, Boston, MA 02115, USA

## Abstract

The choroid plexus (ChP) is a specialized epithelial tissue responsible for production of cerebrospinal fluid (CSF), maintenance of the blood-CSF barrier and regulation of CSF-borne signaling factors. However, despite the ChP’s central role in CSF homeostasis, conserved molecular programs of ChP epithelial function across species and conditions remain incompletely defined. In this study, we integrated eight mouse and human single-cell/nucleus RNA-seq datasets to build a multi-species ChP atlas capturing major epithelial, stromal, vascular, and immune populations. Cross-dataset differential expression identified a conserved epithelial gene program encompassing ion transport, barrier function, ciliogenesis, and secretory signaling, while prioritizing understudied genes including *CLIC6* and *VAT1L*. Co-expression network analysis resolved conserved epithelial meta-programs, and transcription factor regulatory inference identified *OTX2*, *MSX1*, and *GLIS3* among conserved regulators of epithelial gene expression. We also generated a novel Visium HD spatial transcriptomic atlas of mouse fourth-ventricle ChP to orthogonally validate epithelial modules. Lastly, cross-tissue analyses revealed substantial transcriptional overlap between ChP and retinal pigment epithelium, including shared epithelial genes and *OTX2*-associated regulatory relationships. Together, these findings define conserved molecular programs of ChP epithelial specialization, nominate candidate regulators for functional investigation, and provide an integrated cross-species and spatial resource for studying ChP physiology.

## INTRODUCTION

The choroid plexus (ChP) is a specialized epithelial tissue that resides within the cerebral ventricles and forms the blood–cerebrospinal fluid (CSF) barrier. Beyond serving as the principal source of CSF, the ChP performs diverse functions essential for central nervous system homeostasis, including regulating ion and fluid balance, maintaining barrier integrity, transporting nutrients and metabolites, secreting trophic and signaling factors, and providing immune surveillance at the blood–CSF interface(1). Through tightly regulated transepithelial ion transport, metabolic activity, and ciliary function, ChP epithelial cells generate and help circulate approximately 500 mL of CSF daily in adults, maintaining intracranial pressure and supporting neuronal function(2). ChP epithelial junctional complexes and substrate-specific transport systems preserve blood–CSF barrier integrity, while resident immune populations contribute to inflammatory signaling and pathogen defense(3–5). Thus, the ChP operates as a multifunctional secretory organ coordinating cellular programs critical for CNS homeostasis.

Despite its central physiological importance, the molecular architecture that integrates ion transport, barrier maintenance, ciliary motility, and epithelial identity within the ChP remains incompletely defined. Critical elements of CSF secretion, such as Na⁺/K⁺-ATPase activity, bicarbonate transport, carbonic anhydrase function, and aquaporin-mediated water flux, have been studied in isolation, while systems-level regulatory networks that coordinate these processes remain incompletely understood(2, 6). Moreover, dysregulated ChP function contributes to diverse neurological disorders, including hydrocephalus, neuroinflammatory conditions and neurodegenerative disease, underscoring the need to identify clinically tractable molecular nodes governing ChP physiology(1).

Recent advances in single-cell and single-nucleus transcriptomics have transformed our ability to resolve cellular heterogeneity within complex tissues. However, most ChP transcriptomic studies have been limited to single datasets, specific developmental windows, or individual species(4, 7–9). Such studies provide valuable snapshots of ChP molecular physiology within specific biological contexts, but universally conserved ChP transcriptional programs remain undefined. Integrated multi-dataset atlases offer a powerful framework to leverage existing datasets for identification of reproducible transcriptional modules and regulatory circuits across developmental stages, disease states, and organisms(10, 11).

Here, we integrated eight mouse and human single-cell and single-nucleus RNA-sequencing datasets comprising more than 182,000 transcriptomes to construct a cross-species atlas of the mammalian ChP. Cross-dataset differential expression identified a conserved epithelial gene program encompassing transport, barrier, ciliary, and secretory functions and prioritized several comparatively understudied epithelial genes. Co-expression and transcription factor network analyses further identified conserved epithelial meta-programs and regulatory relationships across datasets. We generated a Visium HD spatial transcriptomic atlas of mouse ChP to independently assess the spatial localization and network organization of these conserved programs. Finally, cross-tissue analyses revealed substantial molecular overlap between ChP epithelium and retinal pigment epithelium (RPE), two specialized neuroectoderm-derived epithelia with unusual polarity and transport properties. Together, these analyses define conserved molecular features of ChP epithelial specialization and provide an invaluable resource for future research into ChP physiology.

## RESULTS

### Construction of an integrated multi-species transcriptomic atlas of the choroid plexus

To characterize molecular programs of the choroid plexus epithelium consistent across organisms, developmental time points, and conditions, we integrated eight transcriptomic choroid plexus (ChP) datasets totaling 182,565 transcriptomes across 60 samples(4, 7–9, 12–14) (Fig 1A, Table S1). The integrated atlas included five single-cell datasets and one single-nucleus dataset in mouse, and two single-nucleus datasets in human. Conditions spanned healthy controls, different developmental stages, and various pathologies and pathological models (Fig 1B, Table S1). All major cell types of the ChP were identified including epithelial cells, ependymal cells, immune cells including a myeloid cluster and a mixed neutrophil/lymphocyte cluster, endothelial cells, mesenchymal cells including mural populations and fibroblasts, a mixed neuroglial cluster and proliferating cells (Fig 1C-D). Sub-clustering of vascular/mesenchymal and immune populations further identified traditional cell types within these groups, such as vascular smooth muscle cells, pericytes, T-cells, and dendritic cells among others (Sup Fig 1A-D).

**Figure 1:**
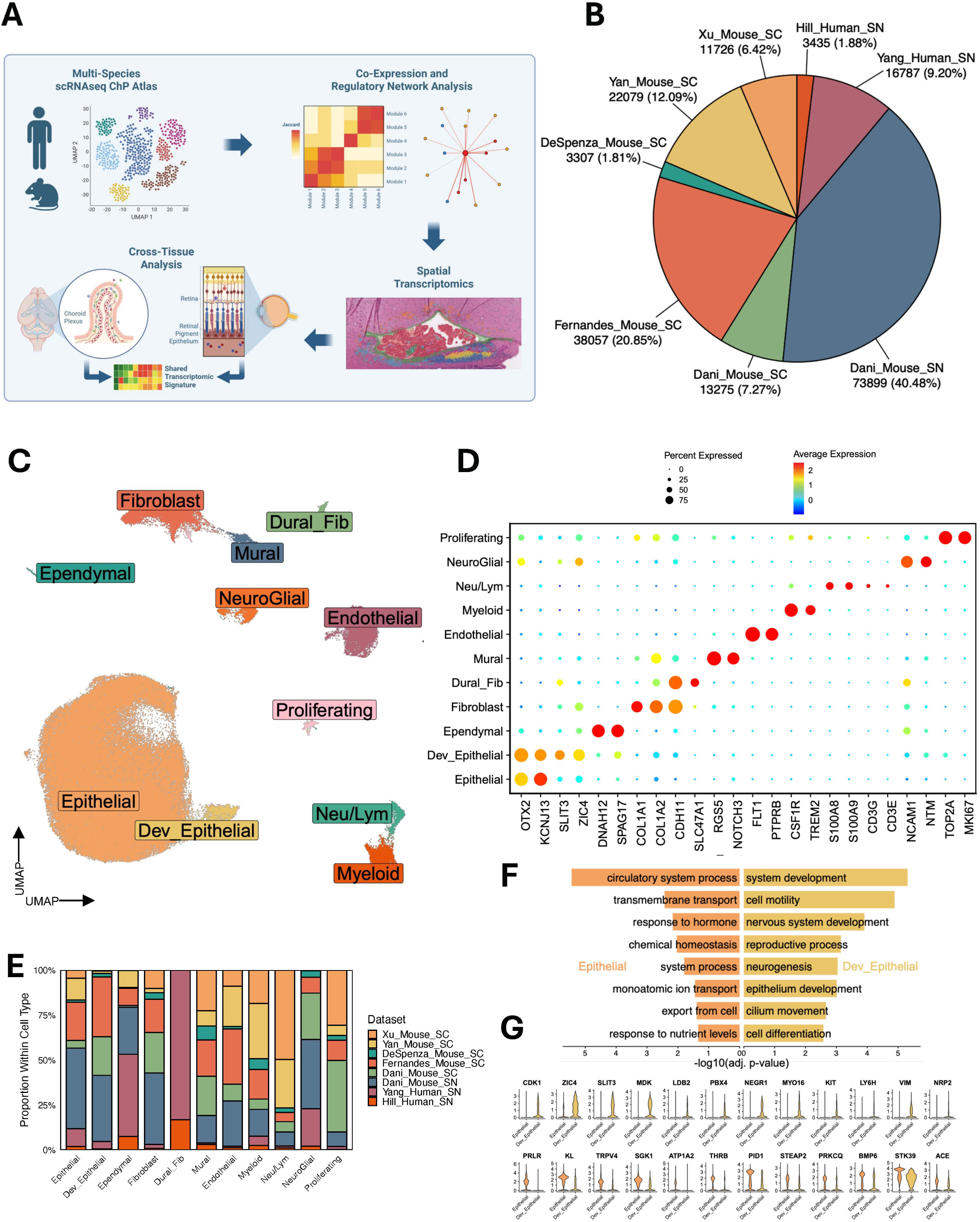
An integrated transcriptomic atlas of the choroid plexus. **(A)** Schematic diagram showing study workflow. **(B)** Pie chart showing number and proportion of cells from each dataset within the atlas after filtering and integration. **(C)** Uniform Manifold Approximation and Projection (UMAP) of all 182,565 integrated transcriptomes from 8 datasets colored by cell type annotation. **(D)** Dot plot showing expression of canonical marker genes across cell types. **(E)** Stacked bar plot showing proportional dataset compositions of each cell type. **(F)** Bar plot showing select enriched Gene Ontology Biological Process terms of differentially expressed genes between Epithelial and Dev_Epithelial cells. **(G)** Violin plots showing expression of select genes driving enrichment of “nervous system development” in Dev_Epithelial cells (top row) and “response to hormone” in Epithelial cells (bottom row).

Importantly, cells from each dataset and both organisms were well represented across clusters, indicating robust cross-dataset integration and relatively homogeneous mixing. The primary exception was a mesenchymal cluster expressing the dural fibroblast marker *SLC47A1*, which was composed exclusively of human samples(15) (Fig 1E, Sup Fig 2A-C). Additionally, among epithelial cells, there was a distinct cluster of developing cells derived from the three datasets of the developing mouse choroid plexus: two datasets of embryonal mouse ChP produced by Dani et al. and a dataset of early postnatal mouse ChP produced by Fernandes et al. (Fig 1C and E). This cluster was enriched for gene ontology terms related to epithelial and nervous system development (Dev_Epithelial) and was characterized by elevated expression of developmental and proliferative markers such as *ZIC4*, *SLIT3*, and *MDK* (Fig 1F and G). In contrast, these cells exhibited reduced expression of canonical transport and hormone-responsive genes, including *PRLR*, *THRB*, and *KL*.

### Identification of conserved epithelial marker genes across datasets highlights canonical choroid plexus function

We next leveraged our integrated atlas to characterize epithelial cells, the most abundant cell type in the ChP. To identify conserved epithelial markers, we performed differential expression analysis comparing epithelial and non-epithelial cells independently within each dataset (Fig. 2A). Unbiased clustering of overlapping differentially expressed genes revealed clear segregation by organism, suggesting species-specific differences in epithelial gene expression programs (Fig. 2B). Despite this separation, 90 genes were consistently upregulated in mouse and human epithelial cells, each exceeding an average log2 fold-change threshold of 1 in each dataset (Fig. 2C, Table S2). These genes were subsequently prioritized based on their aggregate significance across datasets, yielding a consensus ranking of conserved epithelial-enriched transcripts. The chloride-selective ion channel *CLIC6*, a known ChP epithelial cell marker of incompletely characterized function, ranked highest overall followed by *VAT1L*, an gene understudied in the ChP encoding a transmembrane protein often associated with intracellular vesicles. Canonical epithelial markers including *OTX2* and *HTR2C* were also among the top 10 genes, with additional established markers such as *AQP1, FOLR1,* and *ENPP2* similarly represented. Public in situ hybridization data and immunofluorescence staining confirmed epithelial enrichment and characteristic subcellular localization of many of these markers including *AQP1*, *ATP1B1, CLIC6, FOLR1,* and *OTX2* (Sup Fig. 3).

**Figure 2:**
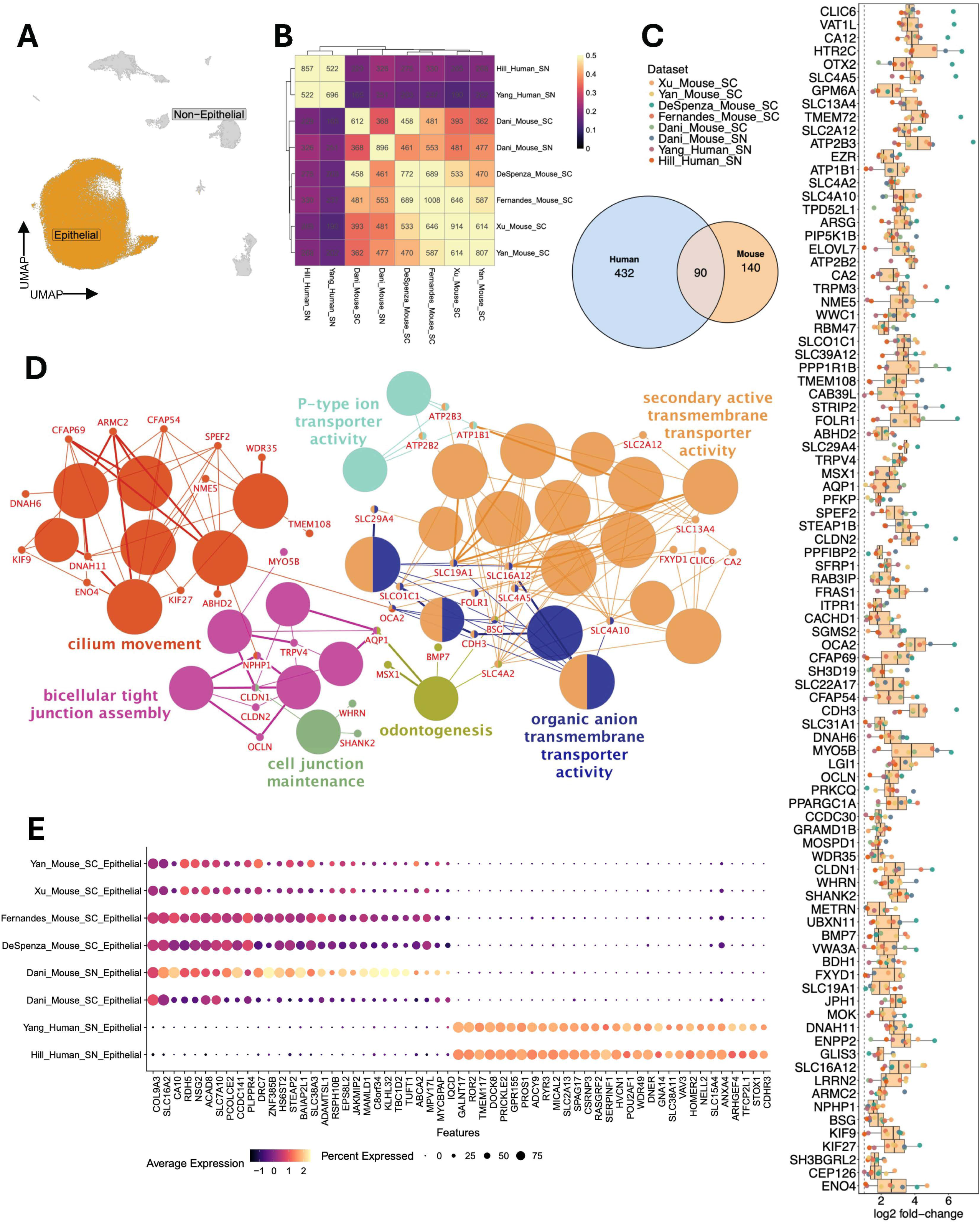
Identification of conserved epithelial marker genes across datasets highlights canonical choroid plexus function. **(A)** UMAP demonstrating epithelial and non-epithelial cell populations used for differential expression analysis. **(B)** Heatmap showing pairwise overlap between datasets with unbiased hierarchical clustering. Fill color indicates the Jaccard similarity index of differentially expressed genes (DEGs) between epithelial and non epithelial cells in each dataset, and the number in each cell denotes the count of overlapping DEGs. **(C)** Bar plot showing distribution of average log2 fold-change values for all 90 genes surpassing an average log2 fold-change of 1 in each dataset. Genes are arranged vertically by mean significance percentile. Inset Venn diagram demonstrates number of DEGs surpassing average log2 fold-change of 1 in all human datasets and mouse datasets. **(D)** ClueGO network of enriched Gene Ontology Biological Process terms of conserved epithelial genes from (C). Large nodes represent GO terms with node size indicating significance and node color indicating ClueGO functional group. The most significant term in each functional group is labeled. Edge weight indicates strength of term-gene relationships. **(E)** Dot plot showing the top 29 mouse-specific and top 30 human-specific genes across datasets.

Gene ontology analysis of conserved epithelial genes revealed strong enrichment for biological processes central to choroid plexus physiology, including terms related to transepithelial ion transport, maintenance of the blood–CSF barrier, and cilia assembly and motility (Fig. 2D). A substantial proportion of conserved genes encoded ion transporters and channels, with multiple members of coordinated transport systems represented. For example, ATPase components essential for establishing ion gradients included Na/K ATPase regulator *FXYD1* and subunit *ATP1B1* along with plasma membrane Ca²⁺-ATPases *ATP2B2* and *ATP2B3*.

Bicarbonate/carbonate transporters (*SLC4A2, SLC4A5, SLC4A10*) ranked among the top conserved genes, together with carbonic anhydrases (*CA2, CA12*) that facilitate rapid CO₂/HCO₃⁻ interconversion. Canonical water channel *AQP1* supported this coordinated fluid-secretion machinery, collectively defining a tightly coupled, energy-dependent secretory cassette. Genes involved in ciliary structure and motility such as *CFAP69, CFAP54* and *DNAH6*, were similarly enriched, along with barrier-associated genes encoding tight junction and adhesion molecules (*CLDN1, CLDN2, OCLN, CDH3*). Conserved transcription factors *OTX2*, *GLIS3* and *MSX1*, together with secreted factors *BMP7*, *SFRP1*, *METRN*, among others, highlighted maintenance of epithelial identity and developmental patterning programs (Sup Fig. 4).

To account for inter-dataset heterogeneity that may obscure conserved epithelial markers under stringent thresholds, we applied a more permissive criterion, requiring an average log2 fold-change greater than 0.5 in both human datasets and in at least five of six mouse datasets. This approach expanded the conserved gene set to 218 transcripts, incorporating additional canonical choroid plexus markers such as *TTR* and *PRLR* (Table S2). Protein–protein interaction analysis of this expanded list revealed four highly interconnected sub-networks highlighting multiple smaller ChP gene relationships (Sup Fig. 5A-E). For example, this expanded network included the gene *STK39* (SPAK), encoding a kinase previously implicated in regulating increased CSF production in inflammatory hydrocephalus(3) and highlighted its relationship to the highly conserved marker and calcium-binding protein *CAB39L* (Sup Fig. 5A). Consistent with our gene ontology findings, these networks were enriched for processes related to membrane transport and barrier integrity, ciliary and axonemal assembly and developmental patterning programs, further delineating coordinated functional modules within the epithelial transcriptome.

Lastly, to assess species-specific differences in epithelial gene expression, we applied stringent filtering criteria to identify transcripts consistently upregulated across all datasets from one species, but not detected above threshold in datasets from the other (see Methods). This analysis revealed 132 genes specific to human datasets and 29 genes specific to mouse datasets (Fig 2E, Table S3). Human-specific transcripts were significantly enriched for genes related to transmembrane transport, with two transporters in particular, *SLC30A10* and *SLC30A1*, driving enrichment in zinc detoxification (Table S3, Table S4). Transporters specific to mouse ChPe cells included *SLC38A3*, from the same family as the human-specific gene *SLC38A11*. Note that these findings likely reflect relative differences in expression rather than true species-restricted gene expression, as transcripts detected in only one species may be present at lower levels in the other, or influenced by species-specific technical or biological factors.

### Co-expression modular analysis reveals conserved epithelial transcriptional programs

To further characterize gene co-expression among cell types of the choroid plexus we conducted high-dimensional weighted gene co-expression network analysis (hdWGCNA) independently in each dataset(16). We clustered resultant co-expression modules based on gene overlap and selected intersecting genes in each cluster to generate 13 predominantly cell-type specific meta-programs (Fig. 3A, Table S5). Hub genes were identified for each meta-program by averaging the degree of connectivity for each gene across modules. A principal epithelial-specific meta-program (meta-program 3) emerged that incorporated the core epithelial module from each dataset (Fig. 3B). Consistent with our differential expression analysis results, this meta-module included many of the shared epithelial markers identified in our differential expression analysis, with 40 of the 50 top hub genes belonging to this group (Fig. 3B, Table S5). Indeed, *CLIC6* was the top hub gene in this meta-module, with the priority of hub genes closely resembling the aggregate significance rankings.

**Figure 3:**
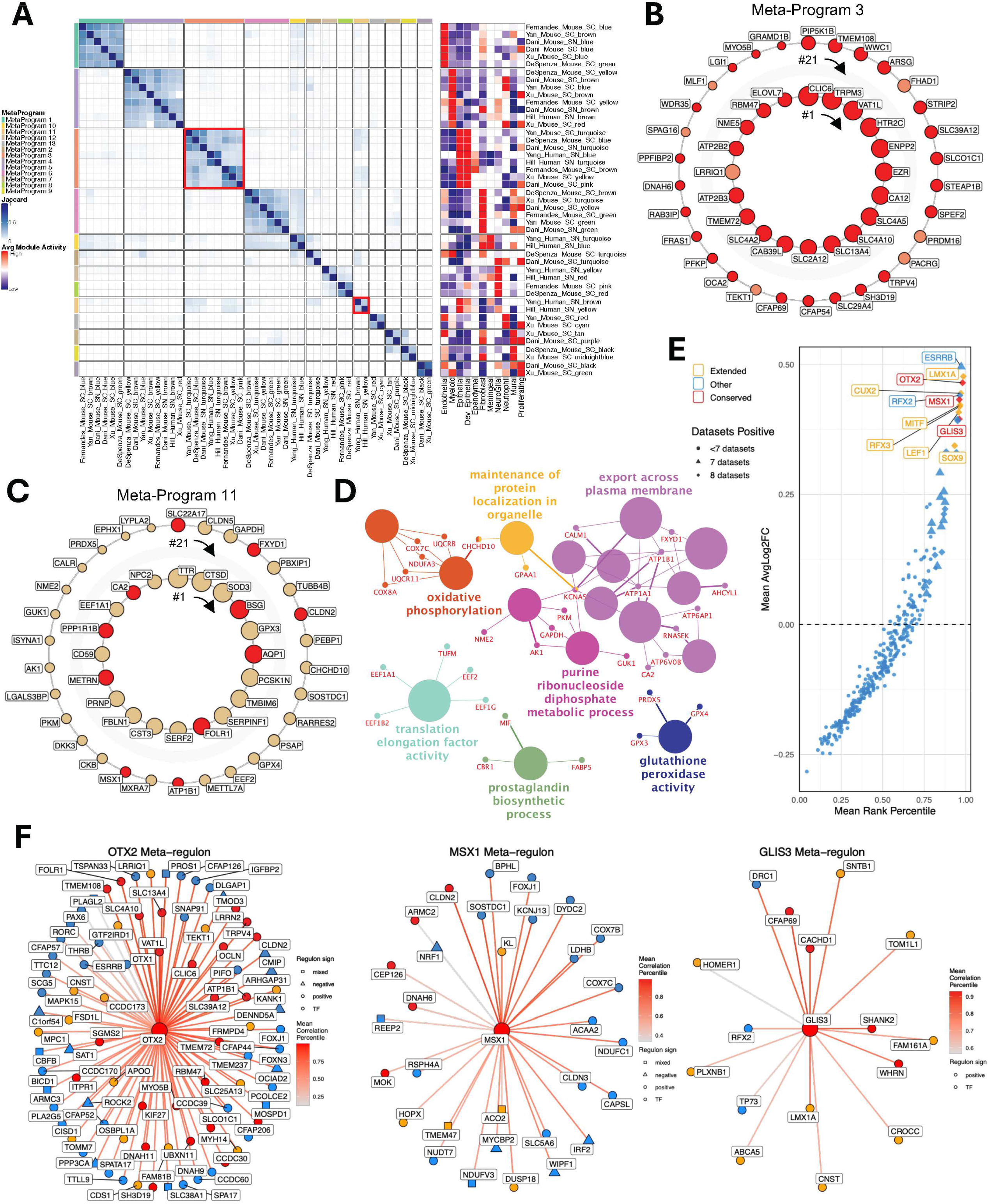
Meta-program and meta-regulon analysis demonstrates shared co-expression and regulatory network relationships across datasets. **(A)** Left: Heatmap showing the Jaccard similarity indices between co expression modules from all datasets. The colored bars along the axes indicate meta program assignments derived from hierarchical clustering of the Jaccard matrix. Right: Heatmap showing the average activity of each module across cell types within cells of the module’s corresponding dataset. **(B)** Circle plot showing top 50 hub genes of Meta-Program 3. Genes are arranged clockwise by connectivity rank with highest rank at 12 o’clock position. The top 20 genes are displayed in the inner ring. Node size indicates gene connectivity rank, and red fill color indicates conserved epithelial gene status. **(C)** Circle plot of top 50 hub genes of Meta-Program 11 as in panel B. **(D)** ClueGO Network of enriched Gene Ontology Biological Process terms of top 100 genes from Meta-Program 11. Large nodes represent GO terms with node size indicating significance and node color indicating ClueGO functional group. The most significant term in each functional group is labeled. Edge weight indicates strength of term-gene relationships. **(E)** Scatter plot showing mean average log2 fold change of regulon scores (Epithelial vs Non-epithelial cells) vs mean regulon rank percentile. Red points are transcription factors from conserved epithelial gene list while orange points are transcription factors from expanded epithelial gene list. **(F)** Regulatory networks of *OTX2*, *MSX1*, and *GLIS3* regulons. Inner, middle, and outer rings represent TF-target gene relationships found across 8, 7, and 6 datasets respectively. Red nodes indicate conserved epithelial gene. Orange nodes indicate genes from expanded epithelial gene list. Circle node shape indicates positive regulation, triangle indicates negative regulation, and square indicates mixed regulation. Edge weight indicates mean correlation percentile.

Notably, a second identified epithelial-specific meta-program (meta-program 11) comprised two modules, each derived from the human ChP datasets (Fig 3A, 3C). Among its top 50 hub genes were the conserved epithelial markers *FOLR1*, *BSG*, *METRN*, *PPP1R1B*, *MSX1*, and *SLC22A17*, together with core CSF secretory machinery components *AQP1*, *CA2*, *ATP1B1* and *FXYD1*. This module also featured the secreted trophic factors *SERPINF1, IGFBP2, METRN* and canonical ChP secreted protein *TTR*, highlighting a potential secretory specialization within this program. Ontological analysis demonstrated significant enrichment for terms related to membrane transport, as well as oxidative phosphorylation driven by inclusion of electron transport chain genes (Fig. 3D). Additional enrichment for glutathione peroxidase activity (*GPX3, GPX4, PRDX5*) and for antioxidant and stress-response genes (such as *SOD3*) suggests coupling of a cellular stress response component within this co-expression module.

Importantly, when meta-modules were constructed separately by organism, nearly all conserved markers assigned to the human-specific meta-module in the integrated analysis were similarly incorporated into the primary epithelial meta-module in mouse (Sup Fig. 6A–B). Indeed, genes such as *BSG*, *FOLR1*, *SLC22A17*, *CA2*, and *FXYD1* emerged among the top 50 hub genes within the mouse epithelial network. These findings indicate that, although these genes were excluded from the primary cross-species epithelial meta-module under stringent intersection criteria, they nonetheless occupy central positions within epithelial co-expression networks across datasets.

### Transcription factor analysis defines regulatory networks of ChP epithelial cells

We next performed transcription factor regulatory network inference independently within each dataset to identify conserved regulons associated with ChP epithelial identity. Several transcription factor regulons consistently ranked among the most differentially enriched in epithelial cells across datasets, including conserved epithelial markers *OTX2*, *MSX1*, and *GLIS3*, along with *CUX2*, *RFX3*, *LMX1A, SOX9, MITF,* and *LEF1* from our expanded conserved marker set (Fig 3E, Sup Fig 7A). To define cross-dataset regulatory architecture, we constructed “meta-regulons” comprising transcription factor–target relationships inferred in six or more datasets (Table S6). Among these, *OTX2* exhibited the most extensive conserved regulatory network, with 11 target genes (including *CLIC6* and *VAT1L*) consistently predicted across all datasets (Fig. 3F). *MSX1* and *GLIS3* similarly demonstrated conserved regulatory influence, with *GLIS3* targeting *LMX1A* across every dataset. Notably, the *GLIS3–LMX1A* interaction appeared bidirectional, and *LEF1* targeted *LMX1A* as well (Sup Fig. 7B). With the additional prediction of OTX2 regulation by *LMX1A,* the meta-regulon suggested a tightly interconnected transcriptional circuit governing epithelial identity (Sup Fig. 7B). Consistent with its established role in ciliogenesis, *RFX3* targeted many cilia-associated structural genes including *KIF27*, and multiple cilia- and flagella-associated protein (CFAP) genes(17) (Sup Fig 7B). These findings collectively define conserved, hierarchically organized transcriptional programs consistently observed in ChP epithelial cells.

### Construction of a spatial transcriptomic atlas validates key conserved nodes of epithelial function

To provide orthogonal validation of findings from our integrated atlas, we generated a Visium HD spatial transcriptomic dataset comprised of three axial sections of the mouse fourth ventricle ChP (Fig. 4A-C, Sup Fig 8A). Spatial spots were subjected to unsupervised clustering with resultant clusters annotated based on canonical marker gene expression (Fig. 4A and B). Across slices, this approach revealed clear anatomical segregation of ChP epithelial cells from ependymal cells lining the ventricular surface and neuronal and oligodendrocyte populations within the surrounding brain parenchyma (Fig. 4C and D). More specifically, in the cerebellum adjacent to the fourth ventricle we observed delineation of the Bergmann glial, Purkinje cell, and granule cell layers (Fig 4C and Sup Fig 8B). Additionally, a small number of vascular stromal cells were distributed adjacent to and within the ChP parenchyma. Importantly, conserved epithelial markers identified in the atlas exhibited robust aggregate activity within spatially resolved ChP epithelium (Sup Fig. 8C).

**Figure 4:**
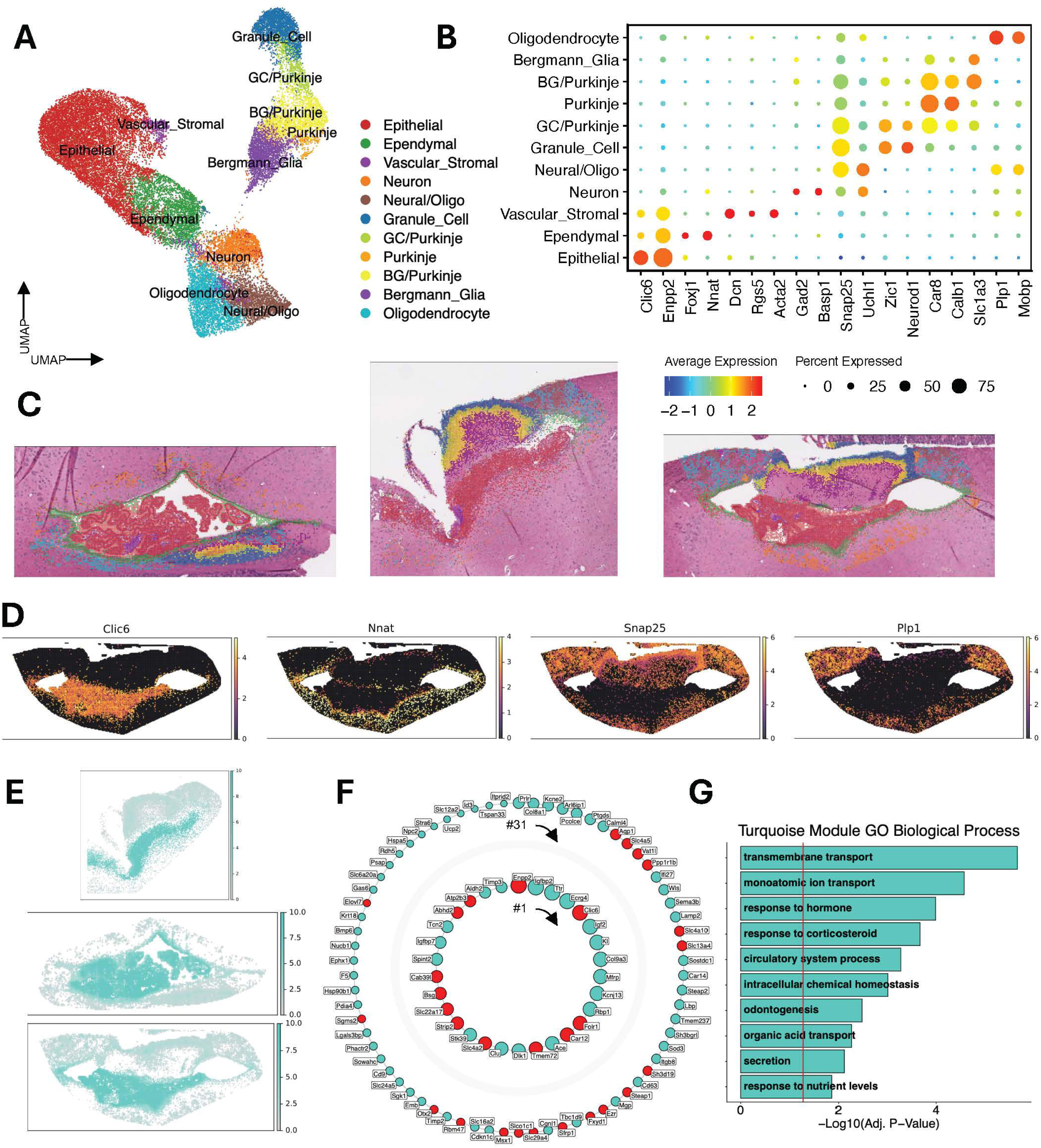
A spatial transcriptomic atlas of the mouse choroid plexus reveals cell-type specific spatial co-expression modules. **(A)** UMAP of spots from Visium HD spatial transcriptomic data colored by cell type annotations. **(B)** Dot plot showing expression of canonical marker genes across spatial cell-types. **(C)** Visium HD sections of mouse choroid plexus colored by annotated cell type. **(D)** Spatial expression of marker genes of epithelial (*Clic6*), ependymal (*Nnat*), oligodendrocytes (*Plp1*) and neural cells in a representative Visium section. **(E)** Spatial activity of the turquoise spatial co-expression module across three representative Visium sections. **(F)** Circle plot showing top 100 hub genes of the turquoise spatial co-expression module. Genes are arranged clockwise by connectivity rank with highest rank at 12 o’clock position. The top 30 genes are displayed in the inner ring. Node size indicates gene connectivity rank, and red fill color indicates conserved epithelial gene status from Figure 2C. **(G)** Bar plot showing select enriched Gene Ontology Biological Process terms of the top 100 genes of the turquoise module.

Spatial co-expression network analysis further resolved 8 modules that were largely cell type–specific and anatomically discrete (Sup Fig 9A-D, Table S7). For example, an ependymal-enriched brown module localized along the ventricular surface and was enriched for cilia and motility-associated genes, while the blue and green modules were respectively specific to neuronal cell types and oligodendrocytes, comprising genes enriched in synaptic function and myelination-related terms (Sup Fig 9A-D). The largest turquoise module was specific to ChP epithelial cells and included 67 of the conserved epithelial markers identified in the atlas (Fig 4E-F). Thirty of these were among the top 100 hub genes, with *Enpp2* ranking highest among conserved markers, followed by *Clic6* and *Folr1*. Consistent with the other mouse datasets, many conserved markers assigned to the human-specific meta-program (i.e. *Aqp1*, *Bsg* and *Fxyd1*) emerged as hub genes within the primary epithelial module (Fig. 4F). Hub genes of this epithelial module were enriched for ontologic terms related to transmembrane transport and secretion (Fig. 4G). Thus, cell populations shared between modalities were characterized by spatialy derived co-expression modules demonstrating strong concordance with atlas-defined meta-programs, providing orthogonal validation of a core epithelial co-expression network.

### Cross-tissue profiling reveals transcriptomic overlap between choroid plexus and retinal pigment epithelium

We next investigated whether transcriptional programs of choroid plexus epithelial cells are recapitulated in other epithelial tissues. Leveraging the epithelial compartment from Tabula Sapiens(18), a human single-cell reference atlas of >1 million cells from 28 tissues, we calculated ChP gene set activity scores across 74 epithelial cell types (Sup Fig. 10A). ChP marker gene activity was highest in ciliated cells of the lung, tracheobronchial tree, and uterus as well as retinal pigment epithelial (RPE) cells (Fig. 5A). Activity in ciliated cell types was driven primarily by the many cilia-related genes within our gene set (Fig. 5B). In contrast, enrichment within RPE cells was driven by a distinct group of transcripts with several ChP genes highly specific to this cell type. This overlapping ChP-RPE signature included many functionally related genes observed in the ChP such as transcription factors *OTX2, SOX9* and *MITF*, secreted morphogens *BMP7, METRN* and *SFRP1*, bicarbonate transporters *SLC4A2, SLC4A5* and *SLC4A10*, plasma membrane Ca^2+^ ATPases *ATP2B2* and *ATP2B3,* and prominent ChP markers *CLIC6* and *VAT1L*, among others (Fig. 5B).

**Figure 5:**
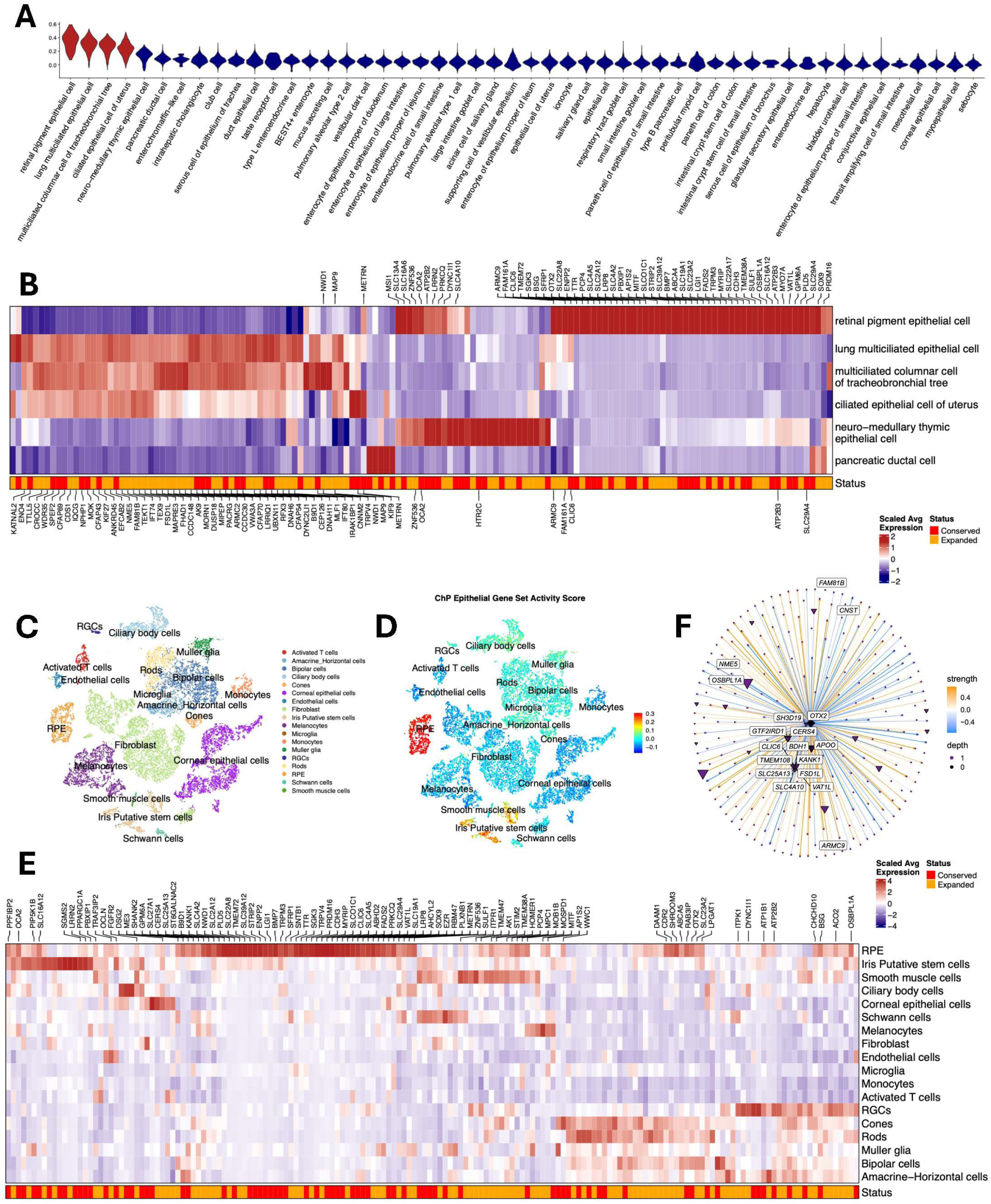
Cross-tissue analysis highlights shared transcriptomic programs of ChP and retinal pigment epithelium. **(A)** Violin plots of expanded epithelial gene set activity score by cell type among top 50 cell types. **(B)** Heatmap of conserved and expanded ChP epithelial genes across top 6 cell types. Top genes expressed in retinal pigment epithelial (RPE) cells labeled on top. Top genes expressed in lung multiciliated epithelial cells labeled on bottom. **(C)** UMAP of cells from transcriptomic dataset of the human eye colored by cell type annotations. **(D)** UMAP of cells from transcriptomic dataset of the human eye colored by expanded epithelial gene set activity score per cell. **(E)** Heatmap of conserved and expanded ChP epithelial genes across cell types. Top genes expressed in RPE cells labeled on top. Only genes with average log2 fold-change greater than 1 in at least 1 cell type are included. **(F)** Regulatory network of *OTX2* regulon. Conserved and expanded ChP epithelial genes are labeled. Edge weights indicate mean correlation strength.

As Tabula Sapiens includes only 67 RPE cells, we conducted identical analyses in a single-cell human eye dataset including 1,011 RPE cells(19) (Fig 5C). ChP epithelial markers were most prominent in RPE cells, with almost identical expression patterns driving this enrichment (Fig 5D and 5E). Regulon analysis demonstrated that *OTX2* up-regulates RPE expression of 17 expanded ChP genes, including *VAT1L* and *CLIC6*, consistent with findings from our ChP *OTX2* meta-regulon (Fig. 5F). Both ChP epithelial and RPE cells are derived from neuroectoderm. The two cell types’ shared reversal of standard epithelial apicobasal polarity supporting specialized ion and fluid transport suggests that their shared transcriptomic signature reflects common mechanisms of secretory function(20, 21).

## DISCUSSION

The ChP epithelium is a specialized interface that plays essential roles in regulation of CSF production and ion composition, secretion of signaling factors and ChP immune function. However, the molecular programs regulating ChP epithelial cell physiology remain incompletely understood. Here, integration of eight mouse and human single-cell and single-nucleus datasets identified a core epithelial gene program reproducibly conserved across species and biological contexts. Independent co-expression and regulatory network analyses converged on many of the same genes, while spatial transcriptomics recovered these conserved components within an anatomically restricted ChP epithelial network. Together, these analyses define conserved molecular features of ChP epithelial specialization and prioritize candidate genes and regulatory relationships for functional investigation.

Atlas construction led to identification of all major cell types of the ChP including developing and mature epithelial cells. Differential expression analysis between developing and mature epithelial cells captured a maturation axis in which developmental/proliferative programs are elevated while transport and hormone-responsive genes are reduced, highlighting maturation dependent transcripts both previously identified (*VIM, KIT, PRLR, KL*) and relatively unappreciated (*LDB2, ZIC4, THRB*)(9). Leveraging the diversity of our atlas, cross-dataset differential expression converged on a compact set of conserved epithelial-enriched genes whose functional enrichment strongly recapitulated known ChP physiology(1). Notably, the two highest-ranked conserved epithelial genes were *CLIC6* (Chloride Intracellular Channel 6), a cancer-associated intracellular chloride channel existing in both soluble and membrane-bound states, and *VAT1L* (Vesicle Amine Transport 1-Like) a transcript predicted to promote oxidoreductase activity and zinc binding recently implicated in Alzheimer’s disease and schizophrenia(22–25). Despite their robust conservation and centrality in our analyses, the functions of *CLIC6* and *VAT1L* in the ChP epithelium have been incompletely characterized, positioning them as high-priority candidates for functional study(26).

Network-based analysis identified a dominant epithelial meta-program derived from all datasets and prioritized many of the markers identified in our differential expression analysis. A second epithelial-specific meta-program derived from human datasets emphasized genes associated with membrane transport, oxidative phosphorylation and antioxidant/stress-response, suggesting an epithelial state reminiscent of ChP epithelial “dark cells”(27, 28). This program contained key nodes involved in CSF production (*AQP1, CA2*), establishment of electrochemical gradients (*FXYD1, ATP1A1, ATP1B1*), and mitochondrial respiration (*COX8A, COX7C*), consistent with a coordinated coupling of bioenergetics to secretory/transport function. Indeed, co-expression of *AQP1*, *SERPINF1*, *CLDN5* and markers of cellular respiration is consistent with “dark cells” characterized in human ChP organoids(29).

Transcription factor regulatory inference elucidated target genes of established regulators of ChP epithelial identity *OTX2* and *MSX1*(30–32) as well as *GLIS3*, a transcription factor with sparse literature directly linking it to the ChP. *GLIS3* is best known for its critical roles in pancreatic β-cell development, thyroid hormone biosynthesis, and kidney function suggesting that it may play a previously underappreciated role in the regulation and development of polarized epithelia specialized for secretion and ion transport(33, 34).

Cross-tissue analysis revealed molecular programs shared between the ChP and other epithelial cell types. Unsurprisingly, ChP cilia-related transcripts were highly expressed in other ciliated cell types of airway and uterus. However, retinal pigment epithelial (RPE) cells exhibited the greatest enrichment of ChP, with highly specific expression of *CLIC6*, *VAT1L* and *GPM6A*, among dozens of other top ChP genes expressed. Epithelial cells of both ChP and RPE exhibit reversed apicobasal polarization, and have been characterized as specialized secretory neural barriers(21, 35). This overlapping transcriptomic architecture of ChP and RPE, including key transcription factor *OTX2*, suggests these genes may play key roles in the unusual polarity and vigorous secretory function of these tissues, promoting several candidates for further study. Indeed, *Clic6*-deficient mice exhibit retinal lamination defects and aberrant microvillar morphology, illustrating how comparative transcriptional analyses can identify transcripts with important roles in tissue identity and function(36).

There are important limitations that should be acknowledged when interpreting these results. First, our cross-study integration necessarily aggregated transcriptomes generated using diverse protocols and the available datasets were not ideally balanced across species or platforms. Thus, to prevent dataset-dependent batch effects from dominating results, we utilized conservative, within-dataset analyses and compared results across datasets rather than relying exclusively on fully pooled analyses. As new publications emerge, future studies may construct atlases with more homogenous datasets, allowing for more integrated analyses across cells.

Second, inclusion of genes in a given gene list or regulon is largely dependent on how inclusion criteria are defined. Thus, we utilized conservative inclusion criteria which may overlook biologically meaningful signals that fall just below detection or vary in baseline expression across datasets. Third, co-expression modules and regulons are based on computational inferences and should be viewed as hypothesis-generating until functionally validated. Finally, spatial transcriptomic data is highly sparse, thus, spatial data suffers from a greater degree of dropout compared to single-cell/nucleus datasets, and a number of spots captured within a section must be excluded due to failing to pass quality control.

In summary, we constructed a multi-species integrated transcriptomic atlas of the choroid plexus and orthogonally validated our findings in a Visium HD spatial transcriptomic atlas of mouse ChP. The repeated prioritization of understudied genes and predicted regulatory relationships provides experimentally testable hypotheses, while molecular convergence with RPE suggests a broader context for investigating specialized neural epithelia. Iportantly, our study provides an invaluable resource for future study of the ChP and an essential reference for comprehensive cell-type expression and spatial visualization of genes of interest in the ChP. Beyond providing an integrated transcriptomic and spatial resource, this study establishes a framework for moving from descriptive ChP atlases toward functional dissection of the conserved molecular programs underlying epithelial physiology.

## METHODS

### Sex as a biological variable

Our integrated atlas included datasets containing both male and female samples. However, sex metadata was not available for some datasets. Thus, no explicit analysis was dedicated to determination of sex as a biological variable. Our spatial atlas included data from both male and females and sex was not considered a biological variable.

### Publicly Available Dataset Curation

The publicly available single-cell and single-nucleus RNA-seq datasets of the choroid plexus used in this study are listed in Supplementary Table 1 and include 2 human single-nucleus datasets in human, 1 single-nucleus mouse dataset in mouse, and 5 single-cell mouse datasets in mouse. The platform, conditions, and ChP tissue ventricle of origin of ChP tissue are listed along with links to each corresponding publications specifying where the data can be obtained.

### sc-RNAseq Processing

Datasets for which unprocessed data was available were downloaded and filtered according to criteria specified in their corresponding manuscripts. Doublets were detected using DoubletFinder(37) (v2.0.4) and removed. Raw counts matrices were extracted from datasets for which only pre-processed data was available. Mouse gene names in mouse datasets were mapped to one-to-one to human orthologs identified via the getBM command from biomaRt(38) (v2.60.1). Additional one-to-one orthologs were identified using the gorth command from gprofiler2(39) (v0.2.3) and by convert_mouse_to_human_symbols from nichenetr(40) (v2.2.0).

sc-RNAseq data was analyzed with Seurat(41) (v5.4.0). Resultant count matrices were loaded into individual Seurat objects and were split by sample. Each object was merged into an aggregate Seurat object so that each sample was contained as its own layer in the RNA assay. Global filtering was applied by removing cells expressing fewer than 500 unique features, and cells with greater than 20% mitochondrial reads in single-cell datasets and greater than 5% in single-nucleus datasets. Features were filtered to include only genes that were included in every dataset. Mitochondrial and ribosomal genes were also filtered.

The count expression values were then normalized with the NormalizeData command. 2500 variable features were identified using FindVariableFeatures and scaled by sample using the ScaleData command with split.by = “Sample”. After running PCA, layers of the Seurat object were integrated using the IntegrateLayers command with method = ‘HarmonyIntegration’ and theta = 0.

Broad clusters were identified using the FindClusters function at a resolution of 0.2, visualized using UMAP, and broadly annotated as epithelial, endothelial, mesenchymal, immune, and neuroglial cells based on canonical markers. Each major cell class was subclustered as above (endothelial and mesenchymal cells were subclustered together) at higher resolution to identify low-quality clusters, as designated by high expression of aberrant marker genes of other cell classes or by low number of unique features and total counts expressed overall. After removal of low-quality cells, the whole dataset was re-clustered using the FindClusters function at a resolution of 0.5, with resulting clusters annotated based on canonical markers.

### Identification and ranking of conserved marker genes of epithelial function across datasets and species-specific marker genes

The 90 conserved markers of choroid plexus epithelial cells were identified by first sub-setting the atlas to each individual dataset. Within each dataset, differentially expressed genes in Epithelial and Dev_Epithelial cells compared to all non-epithelial cells were identified using the FindMarkers command from Seurat with a minimum percent threshold of 0.15. Genes that surpassed an average log2 fold change threshold of 1.0 in every dataset were considered conserved across datasets and species. The expanded list of 218 conserved marker genes was defined as genes surpassing an average log2 fold change threshold of 0.5 in both human datasets and in at least 5 out of the 6 mouse datasets. Genes were then ranked within each dataset by adjusted p-value from smallest to largest and the ranks converted to within dataset percentile scores. For each gene, we then computed a cumulative rank score by averaging the adjusted p-value rank percentiles across datasets.

Species-specific genes were defined as those with a log2 fold change greater than 1.0 in all datasets of a given species and less than 0.25 in all datasets of the opposite species. This gene set was further refined by excluding genes with an average normalized expression greater than 0.5 in any dataset of the opposite species.

### Gene Ontology Enrichment and Network Creation

Gene set enrichment networks were generated with ClueGO(42) (v2.5.10) in Cytoscape(43) (v3.10.2). Ontology was set to Go Biological Process. GO Term Fusion was selected, and the GO tree interval was 3 to 8 with a minimum number of genes per term of 3, and a Kappa score of 0.4. A minimum percentage of genes per term of 3% and 4% were used for shared genes and meta-program 11 genes respectively. A right sided hypergeometric enrichment was utilized, and p-values were corrected using the Benjamini-Hochberg method.

### Protein-protein interaction analysis and clustering

Protein-protein interactions of the expanded list of 218 conserved epithelial marker genes were queried from the STRING database(44) using the stringApp (v2.1.1) within Cytoscape (version 3.10.2), with a confidence cutoff of 0.4. The network was then clustered using the Markov Clustering (MCL) algorithm from clusterMaker2(45) (v2.3.4) with a granularity parameter of 2.5. Gene Ontology (GO) enrichment analysis for each cluster was conducted by retrieving group-wise functional enrichment in the stringApp using a whole-genome background. All p-values were adjusted for multiple comparisons using the false discovery rate (FDR) method.

### Generation of Co-expression Meta-Programs

To create gene co-expression meta-programs we conducted high-dimensional weighted gene co-expression network analysis (hdWGCNA) independently in each dataset using the standard pipeline from the hdWGCNA(16) package (v0.4.08). For each dataset, only cell types with at least 50 cells and genes expressed in at least 5% of cells in at least one cell type were included. Meta cells were generated with the MetacellsByGroups command from cells grouped by cell type and condition, using a nearest-neighbor parameter k = 30 and a max_shared parameter of 15. The co-expression network was generated using the command ConstructNetwork with default parameters and the soft_power selected based on the TestSoftPowers command. The module activity for each cell as defined by the module eigengene was calculated using the ModuleEigengenes command while the eigengene-based connectivity (kME) between each gene and each module was calculated using the ModuleConnectivity command.

After generating co-expression modules in each dataset, we then quantified pairwise similarity between all modules from all datasets using the Jaccard similarity index. The resultant similarity matrix was then hierarchically clustered using average linkage and the dendrogram was cut at a height of 0.85 to define meta-programs of clustered modules. Any singleton modules were dropped from downstream analysis. We then defined each meta-program gene set as the intersecting genes of each module within a meta-program, collapsing any modules from the same dataset into one to not exclude all genes. To calculate meta-program hub gene rankings, the genes within each individual module were ranked by kME value and assigned a percentile score. The kME percentile scores for each intersecting gene in a meta-program were averaged across modules belonging to that program. Meta-program genes were then ranked by their resultant mean kME percentile score.

### Generation of Transcription Factor Meta-Regulons

Transcription factor (TF) meta-regulons were generated by conducting TF regulatory network analysis independently in each dataset using the standard pipeline from the hdWGCNA package (v0.4.08). TF binding motifs within gene promoter regions were identified using the MotifScan command and the JASPAR 2020 motif database(46). The network of TFs and their putative targets was generated using the ConstructTFNetwork command with the standard model parameters from the hdWGCNA pipeline. Regulons were defined by using the AssignTFRegulons command with strategy = “A” (where each gene is assigned to the regulon of its top n TFs), reg_thresh = 0.01, and n_tfs = 10. Positive and negative regulon scores were then calculated for each cell using the RegulonScores command and differential regulon expression between epithelial and non-epithelial cells was calculated using the FindDifferentialRegulons command.

For each dataset we ranked the targets of a given TF by the absolute value of the TF–target correlation and assigned percentile scores. TF-target gene percentiles were then averaged across datasets to generate a mean Pearson correlation percentile score. Relationships were classified as positive or negative if the correlation was consistently > 0 or < 0 across datasets. Otherwise, the relationship was classified as mixed. Meta-regulons for each TF were then defined as TF-target gene relationships identified in at least six of eight datasets.

### Animals

All spatial transcriptomic experiments were performed on wild-type (WT) C57BL/6J mice (founders from The Jackson Laboratory) bred in-house at the Massachusetts Institute of Technology. Tissue was collected from 3 animals, two males and one female, harvested at either 6 months (n = 1) or 18 months (n = 2). Mice received no experimental treatments, injections, dietary modifications, or other manipulations prior to tissue harvest.

All animal experiments adhered to NIH guidelines and were supervised by the Massachusetts Institute of Technology Institutional Animal Care and Use Committee (IACUC) under protocol number 2310000586. Mice were housed under temperature and humidity conditions consistent with the ILAR Guide for the Care and Use of Laboratory Animals.

### Tissue collection and fixation

Mice were deeply anesthetized with sodium pentobarbital (Fatal-Plus, Vortech Pharmaceuticals; 150 mg/kg, i.p.). Adequate depth of anesthesia was confirmed by absence of pedal withdrawal and corneal reflexes prior to thoracotomy. While cardiac activity was maintained, animals underwent terminal transcardial perfusion with ice-cold PBS followed by 4% paraformaldehyde (PFA) in sodium carbonate/bicarbonate buffer, pH 10. Brains were then dissected and post-fixed in 4% PFA, pH 10, overnight at 4 °C on an orbital shaker. Following post-fixation, tissue was washed in PBS. The pH 10 fixative used for both perfusion and post-fixation was prepared by combining 12.5 mL 32% (w/v) aqueous paraformaldehyde, 5 mL 1 M sodium carbonate, and 5 mL 1 M sodium bicarbonate, bringing total volume to 100 mL with deionized water to yield final solution of 4% PFA in 50 mM sodium carbonate/50 mM sodium bicarbonate.

Paraffin processing, embedding, and sectioning were performed by the Tissue Processing & Spatial Imaging Core at MIT for downstream immunohistochemistry and Visium HD spatial gene expression.

### Visium HD FFPE Tissue Preparation and Spatial Gene Expression

FFPE tissue blocks were hydrated by submersion in an ice bath and sectioned on the microtome. 5um thick sections were placed to float on the surface of a 42°C water bath. Selected sections were then placed on slides as per protocol’s instructions (CG000684 RevA), left to air dry, incubated for 3 h at 42°C, left to dry overnight at RT, then stored at 4°C until further processing.

After H&E staining and imaging (see below), coverslips were removed by gentle immersion into water-containing beaker and tissue slides were loaded onto Visium HD slide cassettes (PN#1000669). Tissue sections were destained (incubated with 0.1N HCl at 42°C for 15min on thermal cycler and washed 2x with TE buffer) and decrosslinked using the Visium HD Reagent kit (PN#1000668) following the manufacturer’s protocol, CG000685 RevA. The mouse whole transcriptome probe panel (Visium Mouse Transcriptome Probe kit v2, PN#1000667) was added to the tissue sections as a probe hybridization mix for overnight hybridization (16-24 h at 50°C), followed by post-hybridization wash, probe ligation, and post-ligation wash steps per Visium protocol. The Visium HD slide, 6.5mm (PN-1000670) was thawed, washed, and equilibrated and, together with the tissue section slides, were loaded into the Visium CytAssst instrument, in which release of single stranded ligation products from the tissue upon RNase treatment and their capture on the Visium slide take place. The Visium HD slide was then placed in a Visium Cassette (PN#1000669), the captured ligation probes were extended to generate spatially-barcoded ligation products that were then eluted from each sample. Upon a pre-amplification step, dual-indexed probe-based libraries were generated. Libraries passing the QC step were pooled and sequenced on Illumina S4 lane (300 cycles); for sequencing on the Ultima Genomics platform, libraries were modified by addition of specific adaptors per UG dePCR-Compatible Library Amplification Conversion Protocol, then pooled and sequenced on the UG100 platform (116 cycles).

### Imaging

Slides were deparaffinized and stained with Hematoxylin and Eosin (H&E) according to the 10X Genomics protocol CG000684 RevA (steps involved 2 h incubation at 60° C on a thermal cycler, followed by sequential immersions into Xylene 2x 10 min, 100% EtOH 2x 3 min, 96% EtOH 2x 3 min, 70% EtOH 3 min, water 20 sec, Hematoxylin staining at RT for 1min, washing with sequential immersions into water 5x, 15x, 15x, Bluing buffer staining at RT for 1 min, washing with water by immersion 15x, alcoholic Eosin staining at RT for 1 min, washing with water by immersion 10x). Slides were then temporarily coverslipped in 85% Glycerol mounting medium and imaged at 40x magnification on a Leica Aperio AT2 DX slide scanner.

### Visium HD Sequencing Data Processing and Region Annotation

CytAssist and high-resolution H&E brightfield images were captured using the Leica Aperio AT2 DX scanner and registered to fiducial markers using the Loupe Browser Visium Image Alignment tool. Registered CytAssist images, fiducial registration JSON files, high-resolution microscopic images, and Ultima Genomics generated demultiplexed FASTQ files were processed with Space Ranger using a mouse reference genome (mm10-2020-A) and the Visium Human Transcriptome Probe Set (Visium_Mouse_Transcriptome_Probe_Set_v2.0_mm10-2020-A). Unfiltered 8 μm output matrices generated by Space Ranger were converted to H5AD format using custom scripts and the read_visium function from R package bin2cell (v0.3.4). H5AD files were subsequently converted into Seurat spatial objects using the R package schard (v0.0.1) function h5ad2seurat_spatial in Seurat (v5.1). Across capture areas, sequencing yielded a median of ∼257 nUMI per 8-µm bin.

To focus on the choroid plexus and its immediate microenvironment, each spatial slice was subset to the choroid plexus and a margin of surrounding tissue using Napari 0.5.1. Only 8-µm bins located within the predefined annotated region and designated as being within the tissue were retained, thereby preserving adjacent ependymal and neural tissue while excluding more peripheral regions.

### Spatial Transcriptomic Clustering and Downstream Analysis

Downstream analysis of Visium HD spatial transcriptomic data was conducted using Seurat (v5.1.0). Seurat objects for each spatial slice were merged into an aggregate Seurat object, with each slice contained as its own layer in the Count assay. The raw counts were then normalized with NormalizeData and 2000 variable features were identified and scaled using the FindVariableFeatures and ScaleData commands respectively. After running PCA, the layers of the Seurat object were integrated using the IntegrateLayers command with method = ‘HarmonyIntegration’. Clusters were then identified using the FindClusters function with a resolution of 1. Clusters were visualized using the RunUMAP command and broadly annotated as epithelial, neuroglial, ependymal, stromal, or immune cells, based on canonical markers.

These major cell classes were then subclustered as above at higher resolution to identify low-quality clusters designated by high expression of aberrant marker genes of other cell classes, or by low number of unique features and total counts overall. After filtering the identified low-quality cells from the original combined Seurat object, the combined object was split into layers by slice, and the raw counts were re-processed, integrated and clustered at a resolution of 2.

Resultant clusters were then annotated based on expression of canonical marker genes. Visualization of spatial transcriptomic data was conducted in Python using the sc.pl.spatial command from scanpy(47) (v1.10.2). Conserved epithelial gene set scores per cell were calculated with the sc.tl.score_genes command.

### Generation of Spatial Co-expression Modules

Spatial co-expression modules were generated using the standard pipeline for 10X Visium HD data from the hdWGCNA package (v0.4.05). Spatial coordinates of spots were converted to a dimensional reduction slot called “spatial”. Meta-spots were generated with the MetacellsByGroups command from cells grouped by cell type, using the spatial coordinates as the input reduction, a nearest-neighbor parameter of k = 20 and a max_shared parameter of 15. Co-expression modules were generated using the ConstructNetwork command with minModuleSize = 10, mergeCutHeight = 0.1, and soft_power selected based on the TestSoftPowers command. As above, the module activity for each spot as defined by the module eigengene was calculated using the ModuleEigengenes command while the eigengene-based connectivity (kME) between each gene and each module was calculated using the ModuleConnectivity command. Ontological enrichment analysis of each module was conducted using the gost command from the gprofiler2 package(39) v(0.2.3).

### Cross Tissue Analysis

The epithelial compartment from Tabula Sapiens was obtained from the CELLxGENE database(18, 48). The single-cell data set of the human eye was obtained from the Single-Cell Portal under the accession code “SCP1311”(19). For each dataset, cumulative activity of our extended ChP gene list was calculated in each cell using the AddModuleScore command from Seurat and was averaged per cell type. The *OTX2* regulon from the human eye dataset was derived using the transcription factor regulatory analysis pipeline from hdWGCNA as described above(16).

### Statistics

Differential gene expression between cell clusters was assessed using the Wilcoxon Rank Sum test from Seurat’s FindAllMarkers command, with p-values adjusted by Bonferroni correction. For ClueGO ontological analysis, a right sided hypergeometric enrichment was utilized. P-values were corrected by the Benjamini-Hochberg method. Overlap of gene co-expression modules was measured using the Jaccard similarity index. The resultant similarity matrix was hierarchically clustered using average linkage, and the dendrogram was cut at a height of 0.85. Hub gene rankings in meta-programs were calculated by ranking genes in each individual module by kME value and assigning a percentile score. The kME percentile scores for each intersecting gene in a meta-program were averaged across modules belonging to that program.

### Study Approval

All animal experiments adhered to NIH guidelines and were supervised and approved by the Massachusetts Institute of Technology Institutional Animal Care and Use Committee (IACUC) under protocol number 2310000586.

## Data Availability

Previously published datasets analyzed in this study are available through public repositories, as deposited by the authors of the original studies. The processed integrated atlas will be made available on Zenodo upon publication. Raw sequencing reads and the processed Seurat object for mouse spatial transcriptomic data will be available in GEO upon publication.

## AUTHOR CONTRIBUTIONS

Study conception and design: ED and KTK

Single-cell RNASeq analysis: ED

Confocal imaging: QL

Graphic generation: ET and WCD

Generation of spatial transcriptomic data: LFO, MK, and KG

Spatial transcriptomic analysis: ED, ET, SBF, NP

Writing and review of manuscript: ED, ET, SBF, QL, LFO, PQD, BF, NHM, HC, SLA, KTK

Project administration: ED and KTK

Funding acquisition and supervision: KTK, MK, LT

## AI DISCLOSURE

During the preparation of this work, the authors used the artificial intelligence (AI) tools ChatGPT-4 and ChatGPT-5 throughout the years 2025 and 2026 to aid in data analysis and to improve the readability, language, and grammar of the text .The authors assume full responsibility for ensuring the integrity and accuracy of AI output presented in this article.

## CONFLICT OF INTEREST STATEMENT

The authors have declared that no conflict of interest exists.

## FUNDING SUPPORT

This work was supported by the National Institutes of Health (NIH) under grant R01 NS109358 (to KTK) and by the National Institute on Aging of the NIH under award U01 AG077227.

## ACKNOWLEDGEMENTS

We thank members of the Kahle and Kellis labs for their helpful insights, discussions, and support. We thank Vishal Sarsani for his valuable guidance and insights during the early stages of this project.

## Notes

### Competing Interest Statement

The authors have declared no competing interest.

