## Supplemental_Figures for "An integrated multi-omic atlas of the choroid plexus highlights conserved molecular programs of epithelial function"

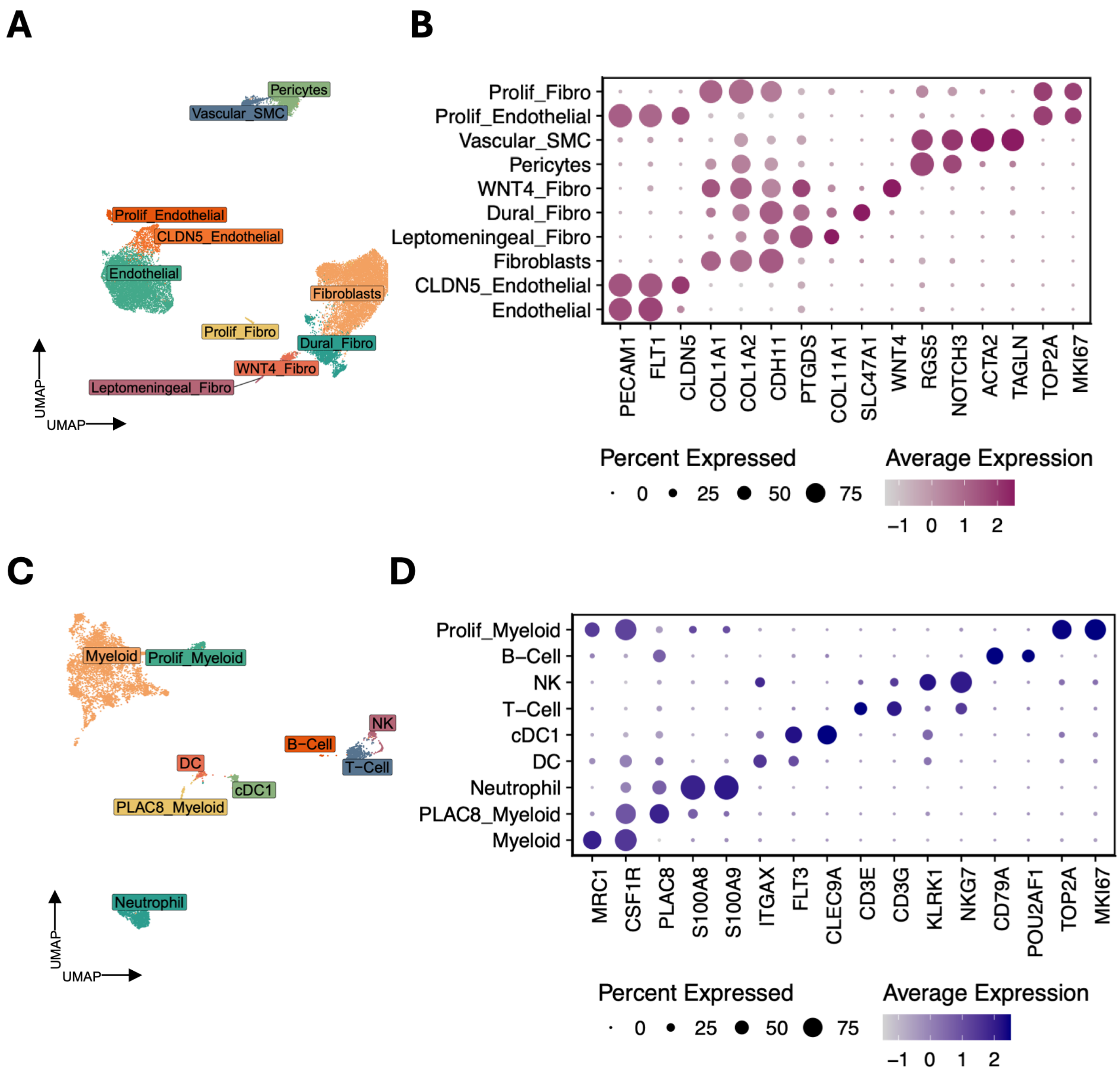


**Supplemental Figure 1: Sub-clustering of vascular/mesenchymal and immune cell populations from the integrated choroid plexus atlas. (A)** UMAP of all vascular/mesenchymal transcriptomes from 8 datasets colored by cell type annotation. **(B)** Dot plot showing expression of canonical marker genes across vascular/mesenchymal cell sub-types. **(C)** UMAP of all immune transcriptomes from 8 datasets colored by cell type annotation. **(D)** Dot plot showing expression of marker genes across immune cell sub-types.

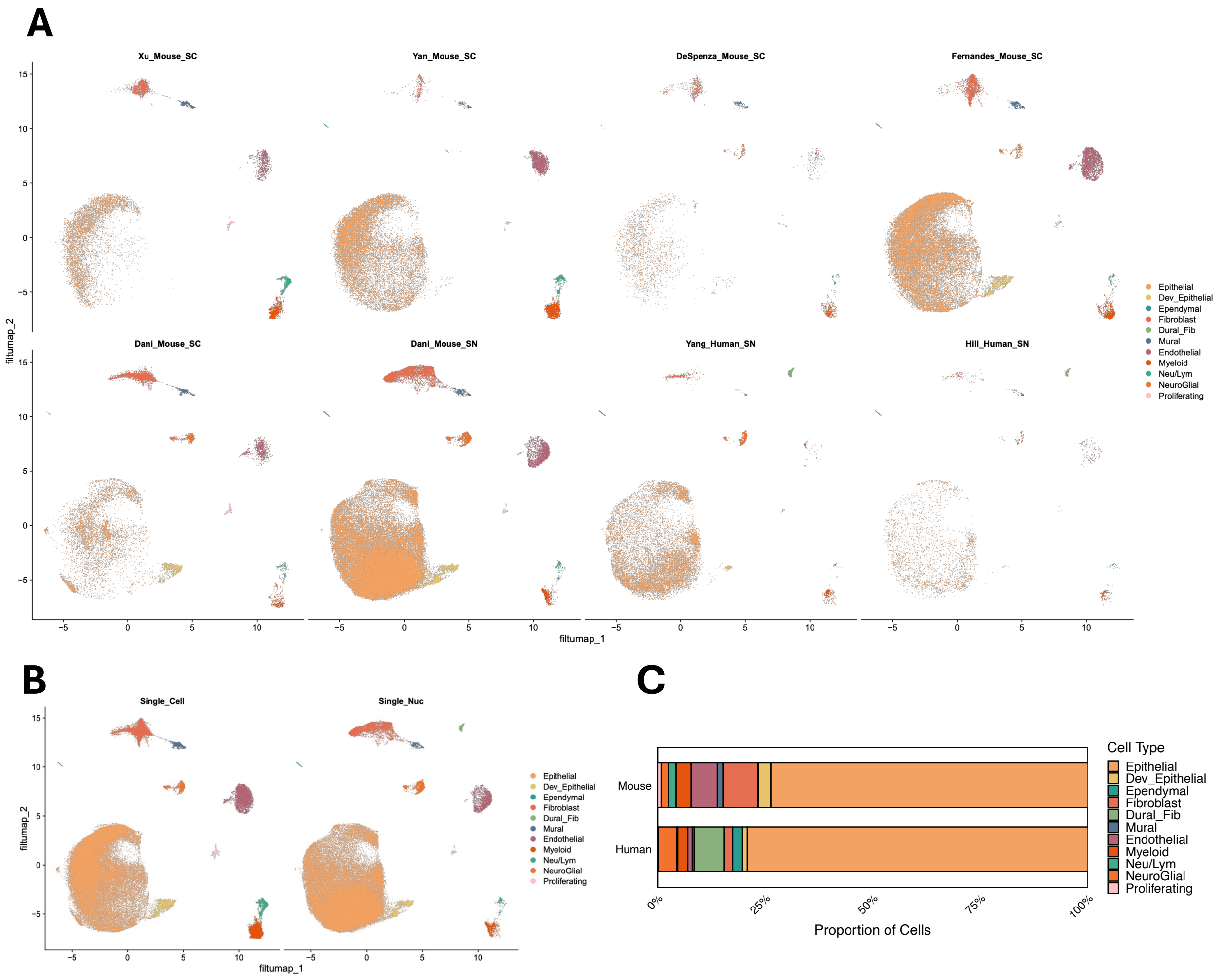


**Supplemental Figure 2: Distribution of cell types from the integrated choroid plexus across dataset, method, and species. (A)** UMAPs of the integrated choroid plexus atlas split by dataset and colored by cell type annotation. **(B)** UMAPs of the integrated choroid plexus atlas split by method (single-cell vs single-nucleus) and colored by cell type annotation. **(C)** Stacked bar plot showing proportional cell type compositions of each species.

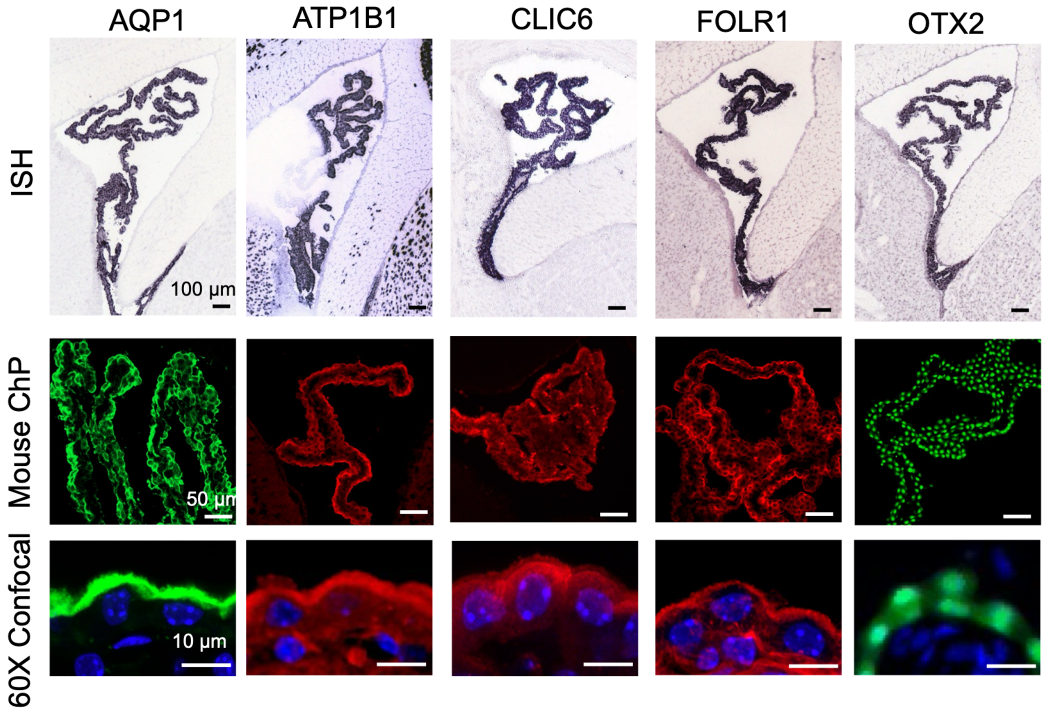


**Supplemental Figure 3:** Representative conserved epithelial markers identified from the cross-species choroid plexus atlas were validated by public in situ hybridization (Allen Brain Atlas) and immunofluorescence in adult mouse choroid plexus. Top, representative ISH images of *Aqp1*, *Atp1b1, Clic6, Folr1,* and *Otx2*. Middle, low-magnification immunofluorescence images. Bottom, high-magnification confocal images showing marker localization in choroid plexus epithelial cells. Nuclei were counterstained with DAPI (blue). Scale bars: 100 µm (ISH), 50 µm (IF), and 10 µm (confocal).

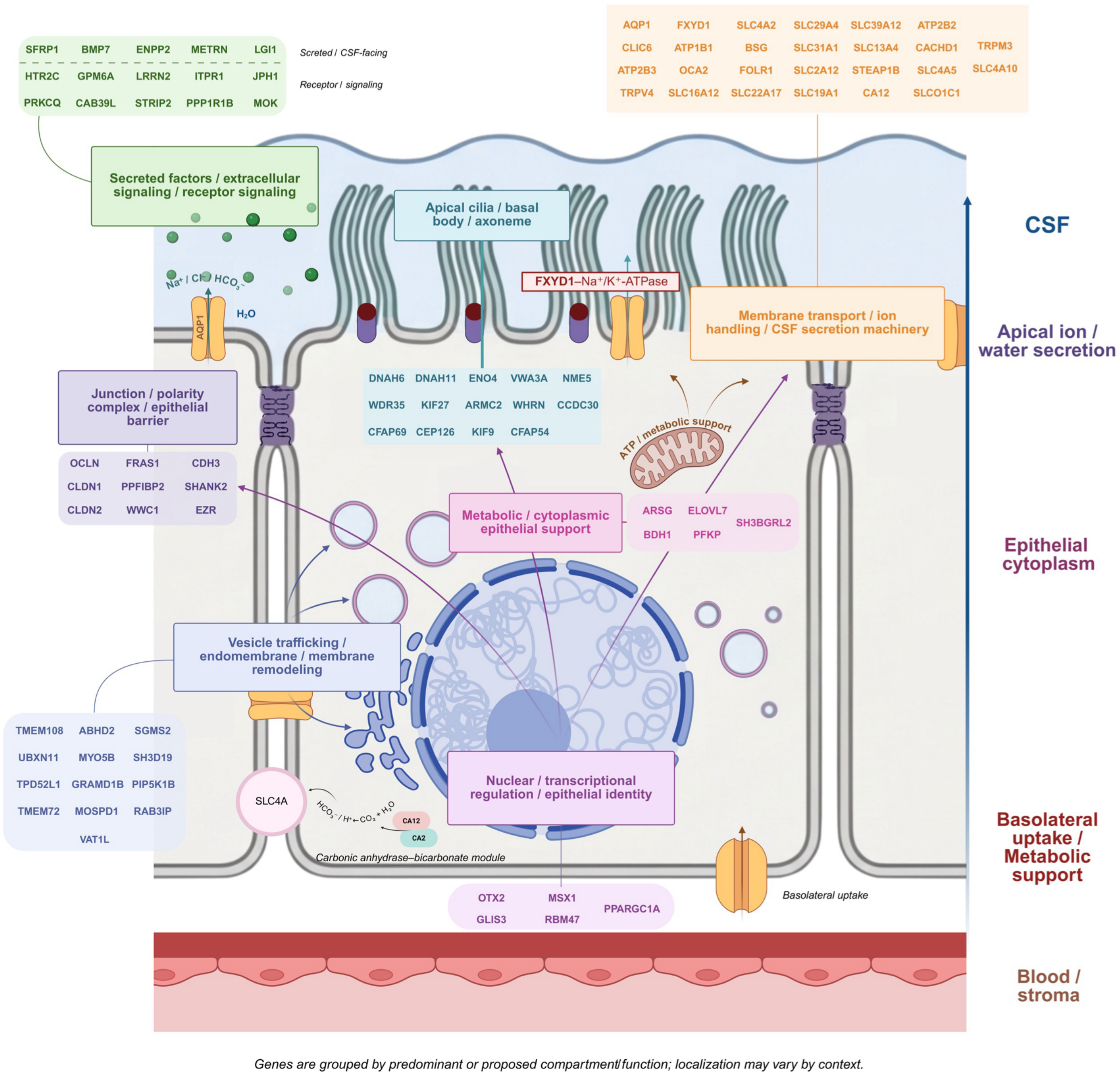


**Supplemental Figure 4:** Biorender graphic depicting cellular location and molecular function of select genes from our conserved epithelial gene list.

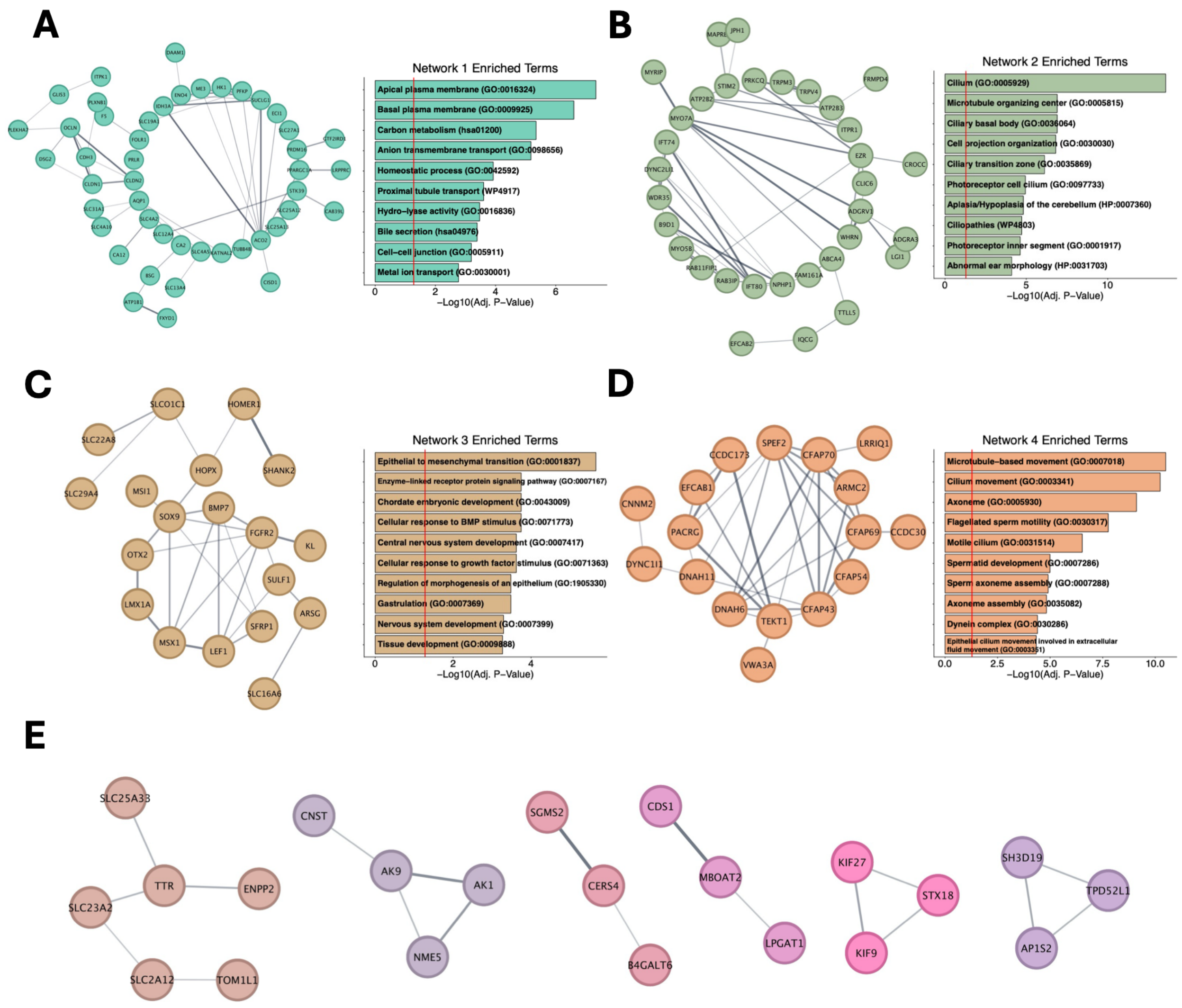


**Supplemental Figure 5: Protein-protein interaction analysis of expanded choroid plexus epithelial gene list. (A-D)** Four largest protein-protein interaction (PPI) sub-networks of the expanded choroid plexus epithelial gene list based on the STRING database (A-D left). Nodes represent proteins; edges represent STRING interaction scores. Bar plots showing top enriched ontologic terms of each sub-network (A-D right). **(E)** Additional PPI sub-networks consisting of at least three proteins.

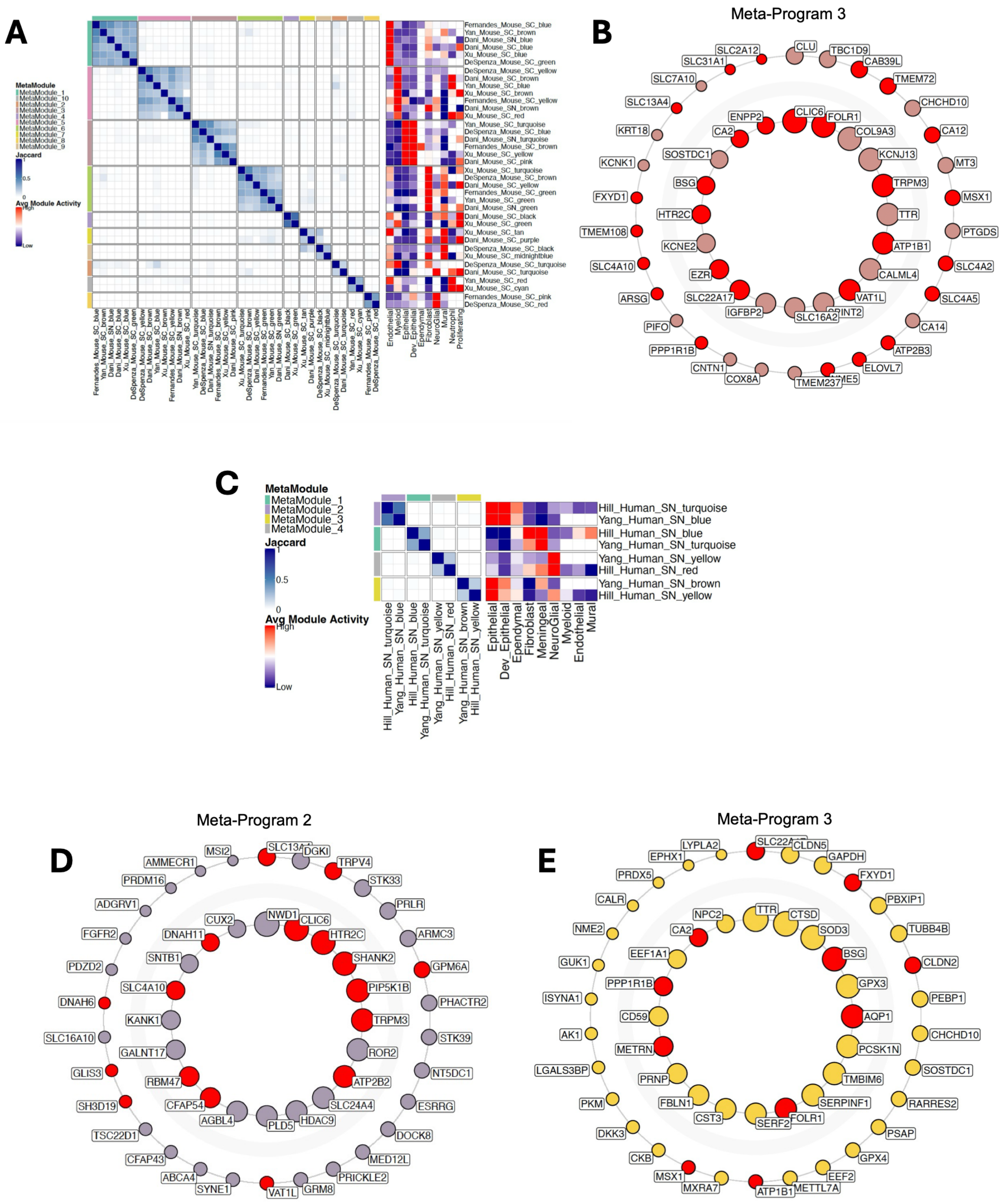


**Supplemental Figure 6: Meta-program analysis of gene co-expression modules by species. (A)** Left: Heatmap showing the Jaccard similarity indices between co-expression modules from mouse datasets. The colored bars along the axes indicate meta-program assignments derived from hierarchical clustering of the Jaccard matrix. Right: Heatmap showing the average activity of each module across cell types within cells of the module's corresponding dataset. **(B)** Circle plot showing top 50 hub genes of mouse Meta-Program 3. Genes are arranged clockwise by connectivity rank with highest rank at 12 o'clock position. The top 20 genes are displayed in the inner ring. Node size indicates gene connectivity rank, and red fill color indicates conserved epithelial gene status. **(C)** Heatmaps as in (A) of modules derived from human datasets. **(D)** Circle plot as in (B) of human Meta-Program 2. **(E)** Circle plot as in (B) of human Meta-Program 3.

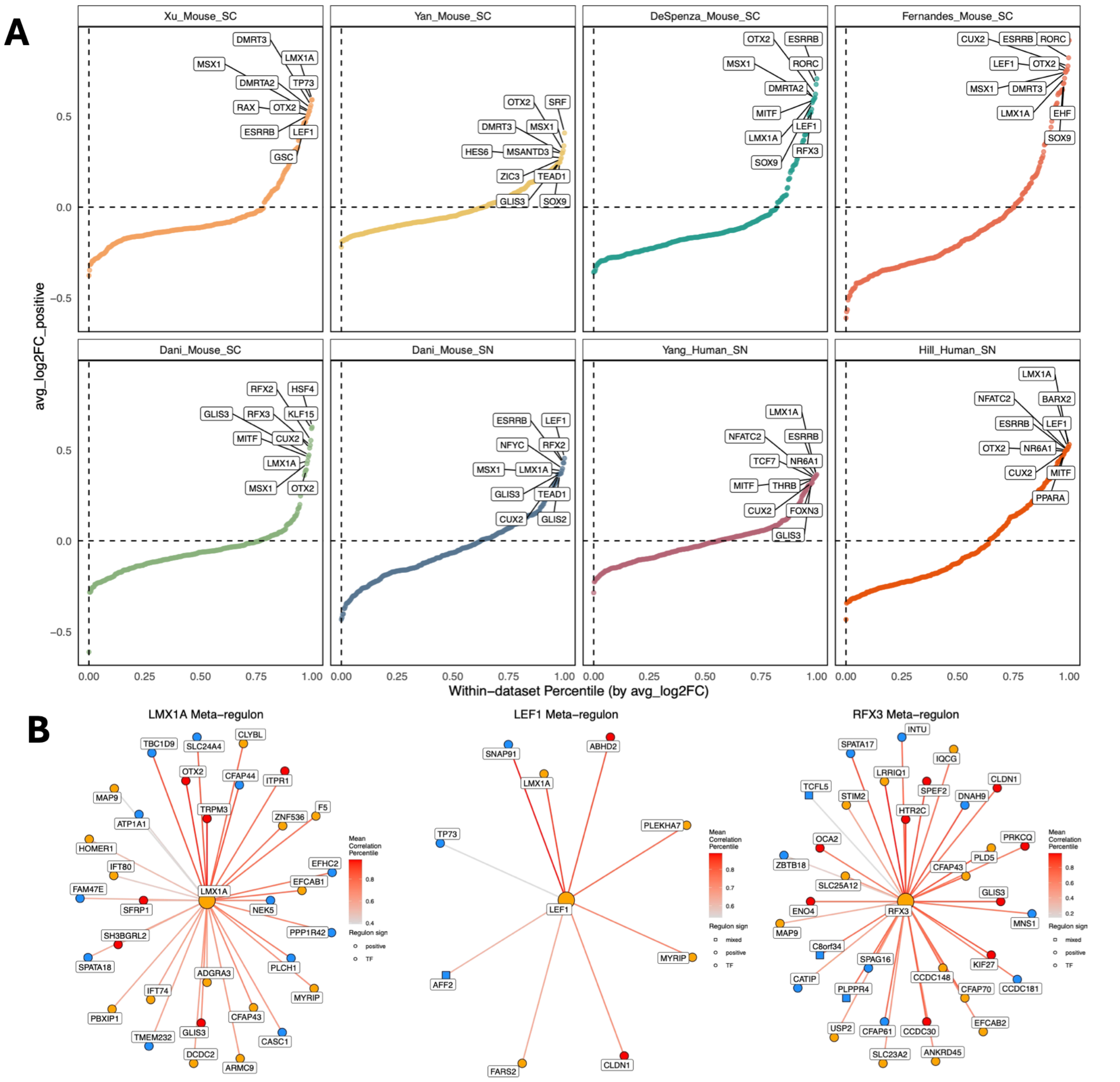


**Supplemental Figure 7: Extended data from meta-regulon analysis. (A)** Rank-ordered log2 fold change plots between epithelial and non-epithelial cells by dataset. The top 10 differential regulons are labeled. **(B)** Regulatory networks of *LMX1A*, *RFX3*, and *LEF1* meta-regulons. Inner, middle, and outer rings represent TF-target gene relationships found across 8, 7, and 6 datasets respectively. Red nodes indicate conserved epithelial gene. Orange nodes indicate genes from expanded epithelial gene list. Circle node shape indicates positive regulation, triangle indicates negative regulation, and square indicates mixed regulation. Edge weight indicates mean correlation percentile.

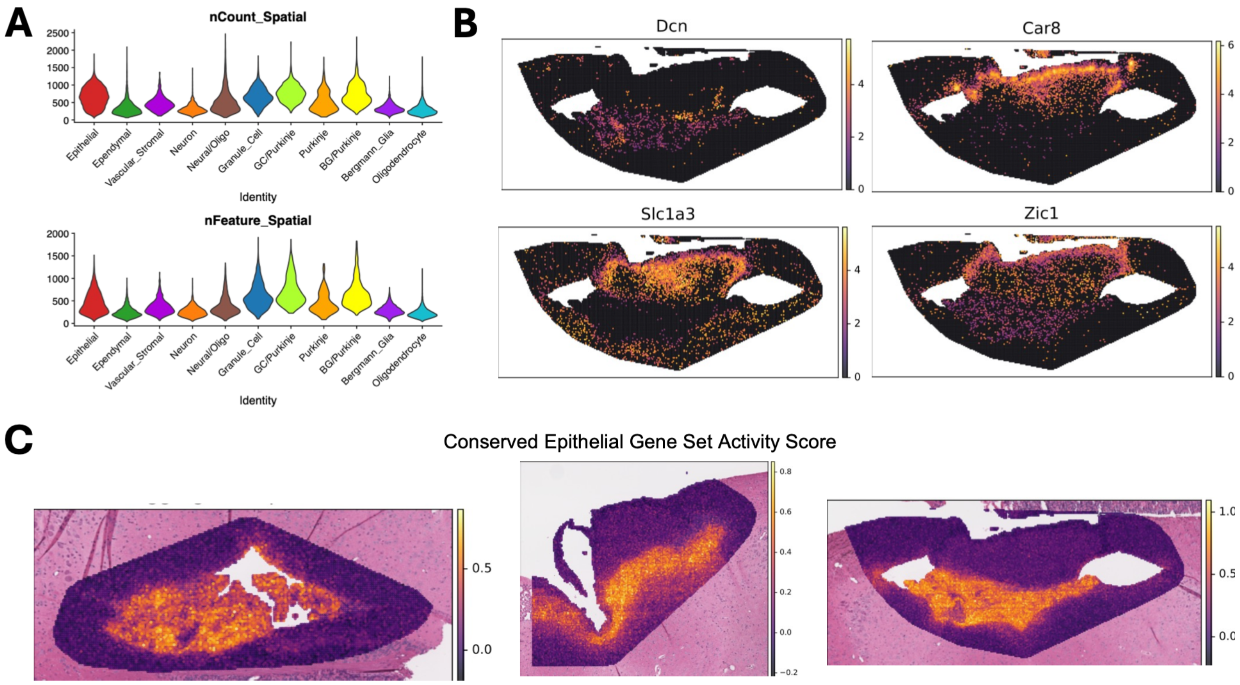


**Supplemental Figure 8: Extended data from Visium HD spatial transcriptomics. (A)** Violin plots showing number of RNA counts and number of unique features per cell type. **(B)** Spatial expression of marker genes of Purkinje cells (*Car8*), Bergmann Glia (*Slc1a3*), granule cells (*Zic1*), and stromal cells (*Dcn*) in a representative Visium section. **(C)** Conserved epithelial gene set activity score per cell in representative Visium sections.

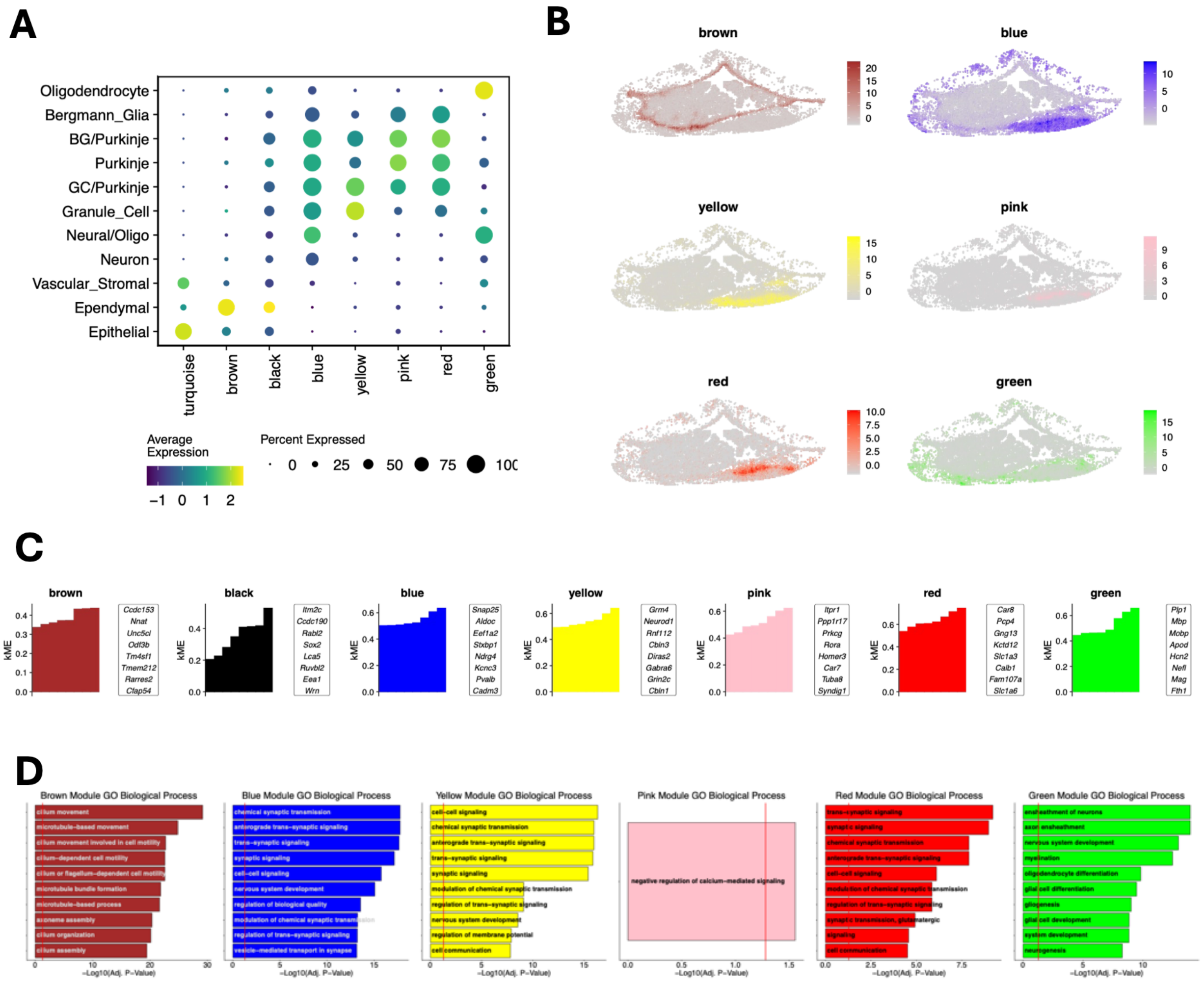


**Supplemental Figure 9: Extended data from spatial co-expression module analysis. (A)** Dot plot showing module activity across cell types. Dot size indicates percent of cells with positive activity score (harmonized module eigengene score) and dot fill indicates average activity score (harmonized module eigengene score). **(B)** Spatial activity of spatial co-expression modules in a representative Visium section. **(C)** Bar plots of eigengene-based connectivity scores of each gene in a module. The top 8 hub genes of each module are listed next to the respective graph. (**D**) Bar plots showing the top significantly enriched Gene Ontology Biological Process terms of each co-expression module.

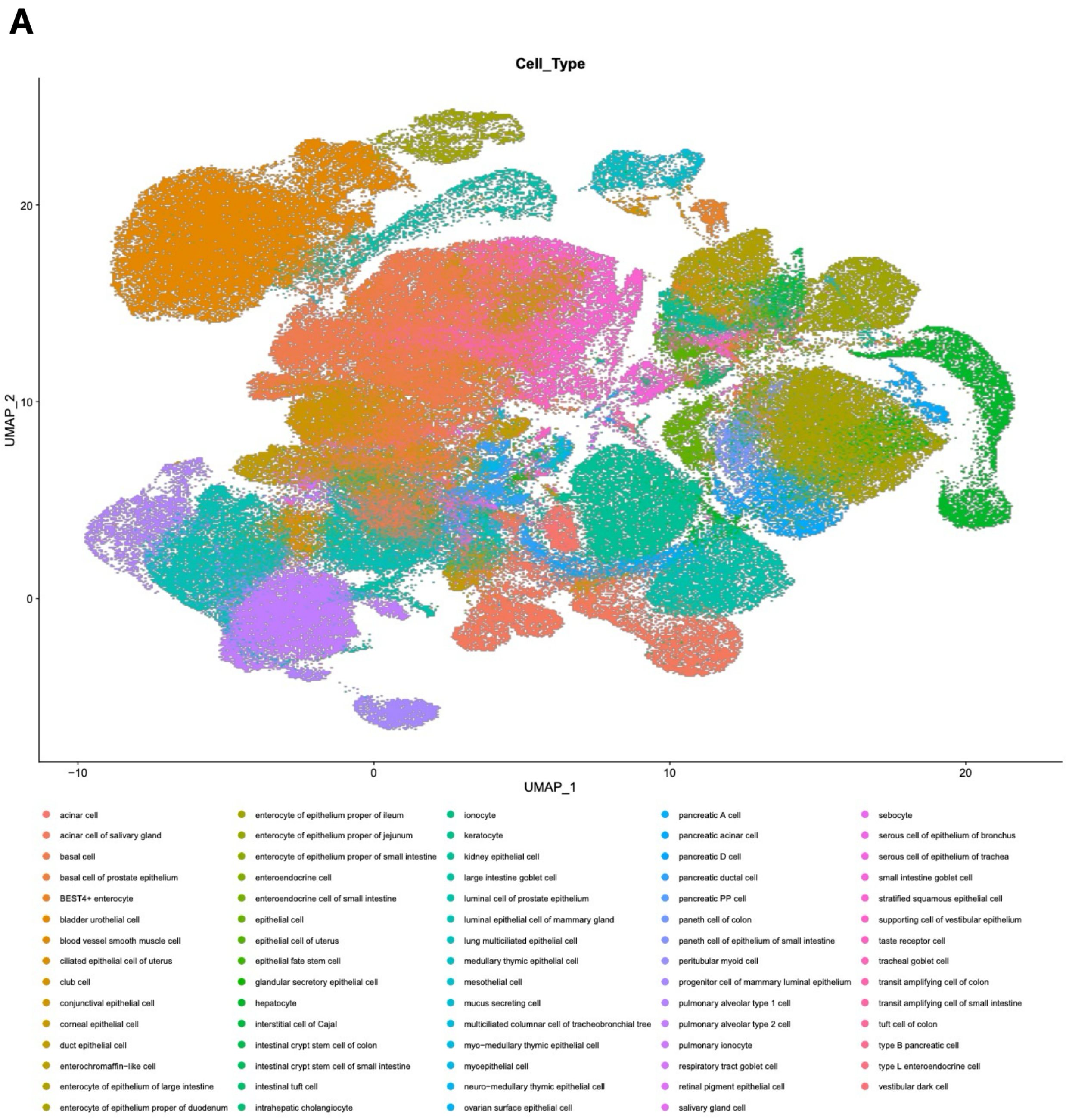


**Supplemental Figure 10: UMAP of Epithelial Compartment of Tabula Sapiens Dataset. (A)** UMAP of epithelial compartment of Tabula Sapiens dataset colored by epithelial cell type
